# A systematic review of post-marital residence patterns in prehistoric hunter-gatherers

**DOI:** 10.1101/057059

**Authors:** Devon D. Brewer

## Abstract

**Background and Rationale:** Post-marital residence patterns refer to where a couple lives after marriage, such as near or with husband’s kin (patrilocality) or wife’s kin (matrilocality). These patterns influence other aspects of social organization and behavior, and potentially reveal key parts of human nature. Since the 1860s, anthropologists have sought to characterize prehistoric hunter-gatherers’ post-marital residence patterns by extrapolating from modern hunter-gatherers’ and chimpanzees’ behavior. For many reasons, these extrapolations are invalid. I summarized direct evidence of residence patterns from prehistoric hunter-gatherers’ remains.

**Methods:** I conducted a systematic review of strontium isotope and mitochondrial DNA (mtDNA) studies of prehistoric hunter-gatherers and extinct hominins. I also carried out a systematic review of the reliability of classifying prehistoric hunter-gatherer individuals by sex. To evaluate assumptions underlying my analyses of residence patterns, I reviewed the ethnographic literature on hunter-gatherers’ mortuary practices as represented in the eHRAF World Cultures database.

**Results:** The archaeologic sites included in my review represent every inhabited continent except Australia, and their dates span almost the last 10,000 years. In the strontium isotope studies, most adults of both sexes were local, and women were slightly more likely to be local than men. Within sites, women and men had similar mtDNA distributions. Women in neighboring, contemporaneous communities had somewhat distinct mtDNA distributions from each other, while the mtDNA distributions of men in the different communities were less distinguishable. The statistical uncertainties for most summaries are fairly large. Taken together, the results indicate the burial communities were mostly endogamous and that exogamous marriages strained toward matrilocality. The very limited research on extinct hominins’ residence patterns is consistent with these findings. In addition, there was only a moderate correspondence between morphologic and genetic estimates of sex in adult prehistoric hunter-gatherer individuals. When I corrected for this measurement error or relied on genetic estimates of sex only, the post-marital residence pattern results shifted in the direction of greater matrilocal tendencies. Modern hunter-gatherers’ burial practices were consonant with the assumptions underlying my analyses.

**Conclusions:** Direct evidence from prehistoric hunter-gatherers’ remains indicates very different post-marital residence patterns than those extrapolated from modern hunter-gatherers’ and chimpanzees’ behavior. In prehistoric hunter-gatherer settings, endogamy may be the outcome of humans’ long-term mating preferences. Endogamy may have inhibited bacterial sexually transmitted diseases in prehistory and enabled the evolution of altruism in humans.

“There is something fascinating about science. One gets such wholesale returns of conjecture out of such a trifling investment of fact.”
– Mark Twain, *Life on the Mississippi*, 1883
“Build up the most ingenious theories and you may be sure of one thing–that fact will defy them all.”
– Fridtjof Nansen, *Farthest North*, 1897

## Introduction

Marriage and kinship are foundational topics in anthropology. In cultures across the world, anthropologists have particularly focused on post-marital residence patterns, or where a couple lives after marriage. In patrilocality (or virilocality), a couple lives near or with husband’s kin. In matrilocality (or uxorilocality), a couple lives near or with wife’s kin. In bilocality, a couple lives near or with either spouse’s kin, perhaps alternating over time. Variations on these patterns also occur. In small-scale societies, these patterns generally involve some sort of exogamy, or marrying a person from another community or social unit. However, when both husband and wife are from the same community, the post-marital residence pattern is called endogamy.

Post-marital residence patterns potentially have far reaching consequences for other aspects of social organization and behavior (1) and may reveal critical parts of human nature. About 150 years ago, early anthropologists such as Bachofen, McLellan, and Morgan asserted that humans in their aboriginal state-presumably hunting and gathering-were matrilocal and matrilineal (tracing descent through the maternal line) (2,3). They based these assertions on ancient classics and ethnographic and archaeologic reports of the period. However, during the early and mid 1900s, the prevailing belief among anthropologists shifted to the view that prehistoric hunter-gatherers were patrilocal (2–4). To support this position, advocates often selectively cited ethnographic reports, including some of suspect integrity (5).

Ember (6) reported the first systematic analysis of the cross-cultural distribution of post-marital residence patterns in modern hunter-gatherers. She found that several different post-marital residence patterns were represented among such cultures in the Human Relations Area Files, and that patrilocality was the most common. Many anthropologists have interpreted Ember’s results and others’ analyses of similar data as indicating that patrilocality was the ancestral residence pattern.

Other anthropologists have challenged these findings by highlighting problems with the specific residence pattern variables and codes in the cross-cultural data sets and the integrity and quality of the corresponding ethnographic data. Marlowe (7) examined post-marital residence patterns by stage of marriage. He found that matrilocality predominated in the early years of marriage, while patrilocality prevailed in the later years. Over the full course of marriage, bilocality was by far the most common. Likewise, Martin and Voorhies (8) showed that the cross-cultural distribution of post-marital residence patterns varied by whether normative prescriptions, acceptable behavior, or observed behavior were used as the measure.

Alvarez (4) demonstrated that the ethnographic data underlying the cross-cultural data sets are mostly very brief statements about normative rules rather than observed behavior. Most statements lacked context and in any case were not based on systematically collected and analyzed data (9). Alvarez (4) also found that genealogical relationships reported by ethnographic informants may not always reflect biological relatedness, as kin categories for some persons can be changed to suit the desires and goals of those involved in making marriages. Furthermore, she noted that some of the post-marital residence patterns codes in the cross-cultural data sets are clearly contradicted by the ethnographies they supposedly represent. Post-marital residence classifications also do not reflect movements of other relevant kin. For example, in some patrilocal societies, wives’ mothers often move to their sons-in-law’s communities (10,11)

Even the ethnographic reports coded for the cross-cultural data sets can belie actual post-marital residence patterns. Dick-Bissonnette (12) presented detailed and extensive evidence on the protohistoric and historic post-marital residence patterns among hunter-gatherer societies in the foothills of the California Sierra Nevada. She showed that anthropologists during the 1900s often attempted to characterize these societies as patrilocal even though the evidence available to them (and also reported subsequently) demonstrated that matrilocality was in fact normative and behaviorally more common. Knight (3) suggested that Engels’ embrace of Bachofen and Morgan’s assertion of ancestral matriliny and matrilocality (and hypothesized matriarchy) left these ideas politically tainted with the rise of communist movements linked to Engels and Marx. Consequently, some anthropologists in western countries during the early to mid 1900s may have been motivated to suppress, distort, or “re-interpret” evidence of matrilocality and matriliny in hunter-gatherers.

Recently, Hill and colleagues (13) analyzed ethnographic censuses from 32 modern hunter-gatherer societies. They found that men were more likely to reside in the same community with consanguineal kin than women were, implying a tendency toward patrilocality. They also observed that in two well-studied societies, most residents of a community were biologically unrelated to each other (from the perspective of any given individual). Hill and colleagues recognized the danger of extrapolating from modern hunter-gather behavior to the ancestral past. Their caution is especially warranted for several reasons. First, few or no modern hunter-gatherers subsist solely from foraging (14–16). Second, modern hunter-gatherers have experienced disrupted subsistence, new ways of storing wealth, new trading and economic relationships, new technologies and goods, geographic displacement, disease, war, and other changes as a result of contact, or even well before contact (e.g., the spread of horses from Europeans in North America to pre-contact societies through intermediaries). Third, and most important, as Murdock (1) underlined, “the one aspect of social structure that is peculiarly vulnerable to external influences is the rule of residence” (p. 201). Post-marital residence patterns, unlike many cultural features, can be changed easily and immediately as the circumstances dictate. For instance, Krech (14,17) and Hanson (18) described post-contact shifts away from pre-contact matrilocal residence patterns in the Great Plains and sub-Arctic of North America.

Some anthropologists and other scientists have also extrapolated patrilocal (male philopatric) residence patterns in chimpanzees backward to the last common ancestor with humans and then forward into human prehistory (for a total of several million years of extrapolation). Koenig and Borries (19) reviewed the genetic, demographic, and behavioral evidence of sex-biased dispersal patterns in the great apes. In most great ape species, both sexes tend to disperse, with females usually dispersing farther. Orangutans (20) and western lowland gorillas (21) are the exceptions, with males tending to disperse farther. Chimpanzees are notable in that only females disperse. Koenig and Borries (19) concluded “… while the genus Pan, particularly common chimpanzees, can be regarded as a good model for human evolution in many respects, its dispersal pattern is not very suitable” (p. 111). Among mammals generally, including monkeys, male dispersal is more common than female dispersal (22,23).

To learn about the past, sometimes there are no alternatives to studying the present for clues. Fortunately, advancing technology in recent decades has enabled direct study of prehistoric post-marital residence patterns. I conducted a systematic review of prehistoric hunter-gathers’ post-marital residence patterns, focusing on two types of analyses of archaeologic human remains.

## Project Overview

I planned this project and specified the protocol in advance. I registered the project at the Open Science Framework (https://osf.io/ea3nm). During the project, I revised the protocol as necessary, as I learned from the literature about the particular kinds of evidence that could be summarized. In nearly every case, I documented my plans in the protocol before conducting any of the additional or revised procedures; later in this section, I report one oversight in my documentation. This article serves as a complete report of all work I did for the project. I have not excluded any activities or results, with one small exception that I describe later in this article.

### Types of studies

I summarized two main types of evidence on prehistoric hunter-gatherers’ post-marital residence patterns:studies of strontium isotope ratios in tooth enamel and studies of the distribution of mitochondrial DNA (mtDNA) variants. I also estimated the reliability of sex estimation for prehistoric hunter-gatherer individuals to put these results in context, and reviewed the ethnographic literature on hunter-gatherers’ mortuary practices to evaluate assumptions underlying my analyses of post-marital residence patterns.

Initially, I had planned to include studies based on phenotypic (cranial or dental) measurements and phylogenetic analyses of cultural evolution as potential evidence on post-marital residence patterns in prehistoric hunter-gatherers. After completing most of the literature search and screening of articles for inclusion in the review (as described below), I decided to exclude such studies. Phenotypic measurements discriminate populations poorly within continents or large regions, and fail the most for geographically adjacent populations (24–26). Moreover, observed variation in adults’ metric cranial measurements did not consistently differ by sex in 9 villages of the modern agro-pastrol Jirel people in Nepal who follow patrilocality (27). In some cases, variation among adults was less in villages with relatively more in-migration. Together, these results demonstrate that comparisons of phenotypic variation by sex are not reliable indicators of post-marital residence patterns, especially on local geographic scales. Many factors affect phenotypic trait expression (28), making it problematic to infer genetic relatedness from similarity of phenotypic traits.

Phylogenetic analyses of cultural evolution in hunter-gatherers involve assessing the relatedness of different modern hunter-gatherer societies (e.g., through linguistic or genetic similarities) and their modern day post-marital residence patterns. Researchers then use phylogenetic analyses to infer the ancestral residence pattern for the proto-society at the root of the phylogenetic tree. I excluded such studies because they are based on ethnographic data on modern hunter-gatherers, which suffer from many methodological problems that prevent reliable extrapolation to prehistoric hunter-gatherers’ behavior, as I noted in the introduction of this article.

### Assumptions

My analyses and summaries of the strontium isotope and mtDNA studies rest on several assumptions about prehistoric hunter-gatherer burial sites:

- Recovered individuals are representative members of their communities.
- A burial site corresponds to a single community over time. For strontium isotope studies, I also assume that the burial site is located geographically within the community’s home range.
- Individuals resided in the burying community when they died.
- Adult individuals were married.

Many of these assumptions are implicit in archaeological and anthropological analyses of prehistoric sites, yet authors rarely state them explicitly. My use of the terms “marriage” and “post-marital” for prehistoric societies is technically incorrect. To my knowledge, there is no direct evidence to indicate that prehistoric hunter-gatherers had practices that would be considered marriage (although it is likely they did). For the purposes of this article, I intend “marriage” to refer to long-term mating in prehistoric peoples.

### Analytic software

I performed the analyses for this project with Gnumeric 1.10.17 (www.gnumeric.org), UCINET 6.504 (www.analytictech.com), Veusz 1.15 (http://home.gna.org/veusz/), and custom programs I wrote in QuickBASIC 4.0 (Microsoft Corporation).

### Structure of methods and results

The four components (strontium isotope studies, reliability of sex estimation, mtDNA studies, and ethnographic accounts of mortuary practices of modern hunter-gatherers) of this project are fairly different from each other in methodological terms. Thus, I present the methods and results in tandem for each component.

## Strontium isotope studies

### Methods

Bentley (29) reviewed strontium isotope methods in archaeology extensively. The ratio of strontium isotopes (^87^Sr/^86^Sr) varies across the terrestrial landscape due to differing geologic substrates and processes as well as atmospheric phenomena, such as precipitation. Consequently, sediments and plants, which take up strontium from the soil, reflect this variation. In turn, animals take up strontium from the plants (and other animals) they eat and water they drink, and deposit it in their bones and teeth. Thus, the strontium isotope ratios in their bones and teeth serve as geochemical signatures of the areas they (or more accurately, the plants and animals they consumed) inhabited. Archaeologists construct local reference ranges of strontium isotope ratios by sampling sediments, waters, plants, and/or animal bones/teeth (both archaeological and modern) in the geographic areas surrounding particular sites.

In humans, bone remodels continually. Therefore, strontium isotope ratios derived from bone samples can reflect strontium deposition over a few to many years. Archaeologic bone samples also may have been infiltrated by ground water, altering the observable isotope ratio in a process called diagenesis. Tooth enamel, however, remains fixed once formed and captures the isotope ratio for the period in which it developed. Enamel is also much less susceptible to diagenesis than bone. Enamel forms on permanent first molars during infancy. On permanent second molars, premolars, incisors, and canines, enamel forms during early to mid childhood. Enamel forms on permanent third molars (wisdom teeth) during late childhood and adolescence (30).

Comparing isotope ratios for an adult individual’s molar(s) with the local reference range indicates whether the individual lived in the local area during childhood and adolescence. Non-local values reflect immigration into the burial community, and thus some kind of exogamy, based on my assumptions. Local values correspond to lifelong residence in the burial community, and thus endogamy. If the burial community had been nomadic, however, strontium isotope ratios might vary considerably between individuals and within individuals (for different molars), and thus yield results that do not indicate post-marital residence patterns.

#### Literature search

I conducted a general literature search of relevant reports, regardless of type of study. I carried out my search in English, but I sought relevant reports in any language and used Google Translate (translate.google.com) to translate abstracts and articles in other languages. To identify potentially relevant reports, I searched Scopus and Google Scholar (scholar.google.com) with the following search terms:(gatherer* OR collector*1 OR forag*) AND (patrilocal* OR matrilocal* OR virilocal* OR uxorilocal*)). Google Scholar returns a maximum of 1,000 records for a single search. To prevent exceeding this limit, I conducted several sub-searches delimited by ranges of publication years. I also used an abbreviated set of search terms (patrilocal* OR matrilocal* OR virilocal* OR uxorilocal*) to search the following additional databases for reports or leads to reports:

- DART-Europe E-theses Portal
- DiVA (Scandinavian theses and dissertations)
- National Electronic Theses and Dissertations Portal (South Africa)
- EThOS (Electronic Theses Online Service) [UK]
- ProQuest Dissertations and Theses (international)
- Theses Canada
- OpenGrey (Europe)
- F1000 Posters
- Figshare
- online abstracts for conferences held by the American Anthropological Association, American Association of Physical Anthropologists, Archaeological Institute of America, Association francaise d‘ethnologie et d’anthropologie, British Association for Biological Anthropology and Osteoarchaeology, Canadian Anthropology Society, European Association of Archaeologists, Finnish Anthropological Association, International Union of Anthropological and Ethnological Sciences, PanAfrican Archaeological Association, Society for American Archaeology, Society for East Asian Archaeology, Society of Africanist Archaeologists, South African Society for Quaternary Research, and World Archaeological Congress (other relevant major professional societies do not appear to have meeting abstracts that are searchable electronically but are otherwise not captured through my other searches)
- Harvard Dataverse Network
- National Institutes of Health RePORTER (grant database, USA)
- National Science Foundation (grant database, USA)

I performed the Scopus, Google Scholar, and other database searches in March and April of 2015. In addition, I examined my personal library, references in reports retrieved from these searches (backward citation search), and sources citing the prior reports (as indicated by Google Scholar; forward citation search) for other relevant reports to include. I completed the backward and forward citation searches between March, 2015, and March, 2016.

After identifying potentially relevant reports, I attempted to retrieve them for detailed examination and abstracting, as appropriate. I retrieved potentially relevant reports from free/open access sources online, through the electronic and physical holdings of the University of Washington libraries, through inter-library loan, and directly from authors themselves. I excluded studies from consideration if they focused on post-contact hunter-gatherers’ remains or involved prehistoric populations with any evidence suggesting even partial horticultural, agricultural, pastoral, or industrial subsistence.

Some reports did not have results sufficient for inclusion in the review, but had information that suggested the authors might have data that could produce results sufficient for inclusion if reported more fully or re-analyzed appropriately. In such cases, I made at least three attempts over a period 4 weeks or longer to contact the corresponding author (and sometimes co-authors) for such studies by email, telephone, fax, and/or post. I requested that the authors perform the particular analyses and share the results, or share the relevant data so I could perform the analyses.

After the initial publication of this article, some authors sent me additional reports. I included two in this revision.

#### Inclusion criteria

I included a strontium isotope report in the review if the sample had at least three adult individuals (including at least one male and one female), the isotope analyses were based on tooth enamel, and the authors estimated a local reference range of isotope ratios. However, I made one exception to these rules:I included one report on isotope analyses of bone from extinct hominins, about whom there is very little empirical evidence. From reports on anatomically modern humans, I included only adult individuals (estimated age > 15 years) in my summaries. Although some individuals just above this age threshold may not have been of marriageable age, many-especially females-undoubtedly were. Any error I introduced by including these very young adults is likely less than that which would be introduced by excluding young married individuals. For sex-specific analyses, I included only adult individuals for whom the authors determined sex confidently (by the authors’ criteria, regardless of their methods of sex determination). I excluded individuals from these analyses whose estimated sexes were “probable.”

#### Data analysis

I performed meta-analyses of the tendency toward endogamy or exogamy and the tendency toward matrilocality or patrilocality. I also reviewed, in narrative form, relevant studies not included in these meta-analyses.

To quantify the extent of endogamy/exogamy, I conducted a random effects meta-analysis (31) of the proportions of individuals (overall and by sex) classified as local. I used the Freeman-Tukey double arcsine transformation (32) to convert observed proportions into a form suitable for random effects meta-analysis. This meta-analytic approach effectively weights each site’s effect size (proportion, in this case) by its sample size when producing a summary estimate. The random effects approach also accounts for observed variation in results as a function of sampling variation as well as genuine differences between studies (sites) in underlying effect sizes. I calculated the I^2^ statistic (33) which indicates the proportion of variation between observed effect sizes that is due to genuine differences in their underlying (“true”) effect sizes. After performing these analyses, I computed back transformations (34) of the summary estimate and 95% confidence limits for the summary estimate and individual sites’ results.

To assess the tendency toward matrilocality or patrilocality, I computed a phi (Pearson) correlation and corresponding 95% confidence interval between individual sex and migration status (local or immigrant, as classified by the authors from the isotope analyses) for each site. I then summarized these correlations with a random effects meta-analysis (31). I displayed individual site and summary results with forest plots.

For sites in which at least three individuals were classified as immigrants who also included at least one individual of each sex, I compared males and females on the extent of the absolute deviation of their isotope ratios from the midpoint of the local reference range. For each of these sites, I computed a point biserial (Pearson) correlation and corresponding 95% confidence interval between individual sex and absolute difference between an individual’s ratio and the midpoint of the local reference range. I performed a random effects meta-analysis to summarize these correlations. Local reference ranges provide reasonable benchmarks for assessing migration, but they are not necessarily definitive, no matter how they are defined. It is impossible to know with certainty the geographic range that a given group of prehistoric hunter-gatherers traversed for obtaining food or the temporal pattern in which they did so. The extent to which isotope ratios deviate from the local range likely correlates positively with distance from the local area. If the local reference range were defined too narrowly, locals could have been misclassified as immigrants, possibly producing little difference between women and men. If that were the case, these analyses would indicate the prevailing post-marital residence tendencies.

### Results

#### Literature search

Figure 1 shows the results of the overall literature search on prehistoric hunter-gatherers’ post-marital residence patterns, including both strontium and mtDNA studies. During the database search stage, the full text of some sources was immediately available, enabling detailed screening with little extra effort. I did not count sources that I screened in this way and excluded as retrieved reports. The summary in figure 5 does not include reports I relied on for demographic data but which do not contain strontium isotope ratio or mtDNA results. A supplementary file (ReportsExcludedFromSystematicReviewOfResidencePatterns.pdf) for this article lists the citations to retrieved reports I excluded from the review and my reasons for excluding them. The count shown in the top left box of Figure 1 represents unduplicated records within, but not between, databases. The count shown in the top middle box in Figure 1 represents the sum of records returned for different forward citation searches in Google Scholar, which are unduplicated within, but not between, searches.

**Figure 1.**
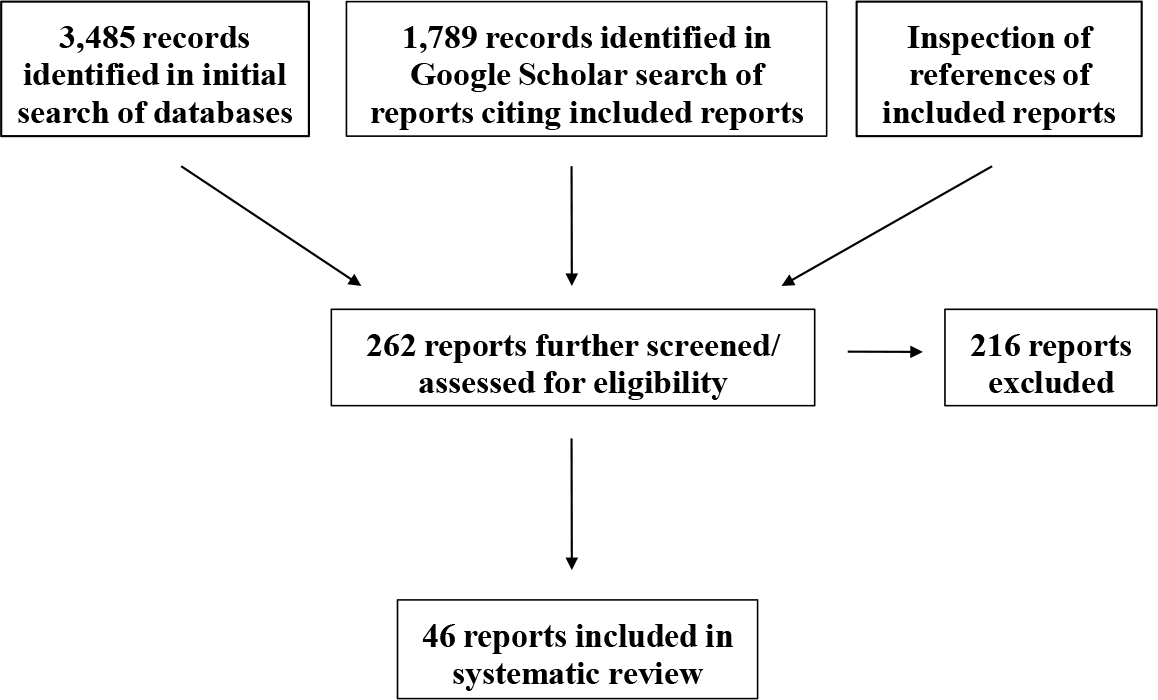
Results of the literature search for strontium isotope and mtDNA studies

Notably, many reports included all the information necessary for my analyses. The vast majority of authors I contacted were helpful and provided the additional data or information I needed for analysis. Other authors were unable to share data because they were no longer available due to death of a colleague or other reasons. However, a few authors never responded to my requests despite many repeated contact attempts and, in a few cases, efforts by intermediaries. Several authors also refused to share such additional information, and some declined to share conference presentations or forthcoming publications that had already been cited or summarized elsewhere. The supplementary file notes the reports that I excluded because I could not get access to the relevant data.

#### Study characteristics

Table 1 shows the key characteristics and results for the eight strontium isotope studies included in the meta-analyses. The sites represent all inhabited continents except Australia. The dates of the sites extend from nearly 10,000 to several hundred years before present. Most authors noted that the sites appeared to represent sedentary, or semi-sedentary (seasonal occupation), communities. None of the authors hypothesized a particular post-marital residence pattern. Indeed, only one author (35) explicitly examined post-marital residence patterns. Nonetheless, nearly all authors reported in their publications the isotope measurements for each individual and other information necessary to assess these patterns. None of the included studies was pre-registered. Except for Quinn and colleagues (36), all study teams received funding support from national government agencies and some also had support from universities and/or private foundations.

#### Meta-analyses

Table 1 also shows that the large majority of both women and men were classified as local at every site except Gobero, indicating a fairly strong tendency toward endogamy (marriage within the community). At two sites, Ota and Tsukumo, all adult individuals were local. The random effects summary estimate across sites is that 83% of prehistoric hunter-gatherer individuals were local. A somewhat higher proportion of women than men were local. Genuine variation in these proportions across sites was considerable, as indicated by large I^2^ values (overall =.87, men =.84, women =.74). The Gobero estimate, in particular, is a prominent outlier.

Figure 2 and the last column of Table 1 show the phi correlations, when calculable, between sex and local/immigrant status. In the five studies in which individuals displayed variation in local/immigrant status and included confidently estimated male and female adults, matrilocal tendencies were present in four. The resulting summary estimate for these studies indicates a weak overall tendency toward matrilocality (weighted mean phi correlation =.09, 95% CI:-.06-.24). Within-study sampling errors could account for all of the observed variation in correlations (I^2^ =.00), meaning that there were no detectable genuine differences between sites. There was no consistent sex difference in deviations from the local strontium ratio range among those individuals that authors classified as immigrants (mean phi correlation = -.09, 95% CI:−.36 −.20, I^2^ =.00, *n* = 4 studies; Table 2). This means that any error or bias in the authors’ definitions of the local isotope ratio range probably did not affect the assessment of post-marital residence patterns significantly.

**Table 1.**
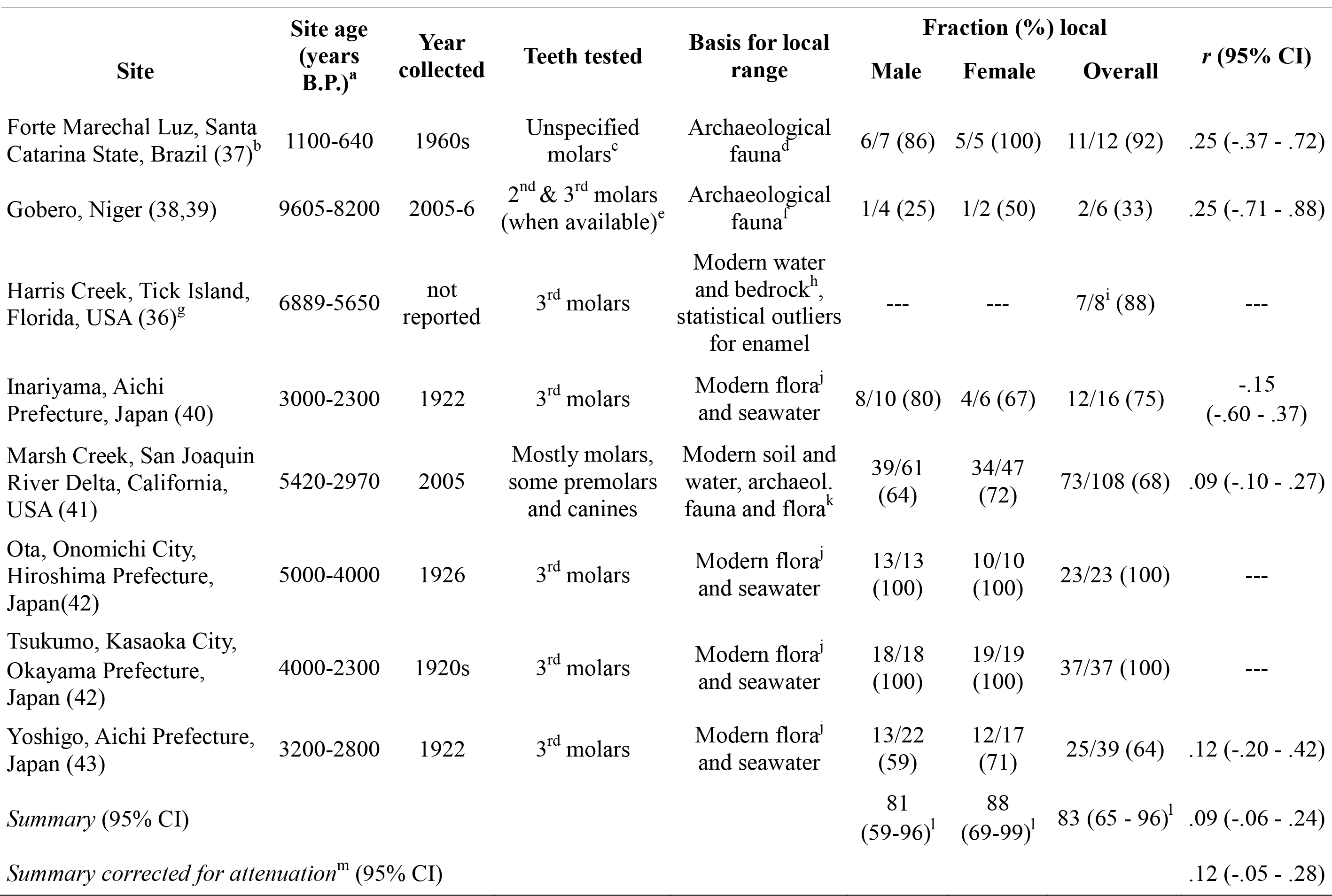
Strontium isotope studies of prehistoric hunter-gatherers

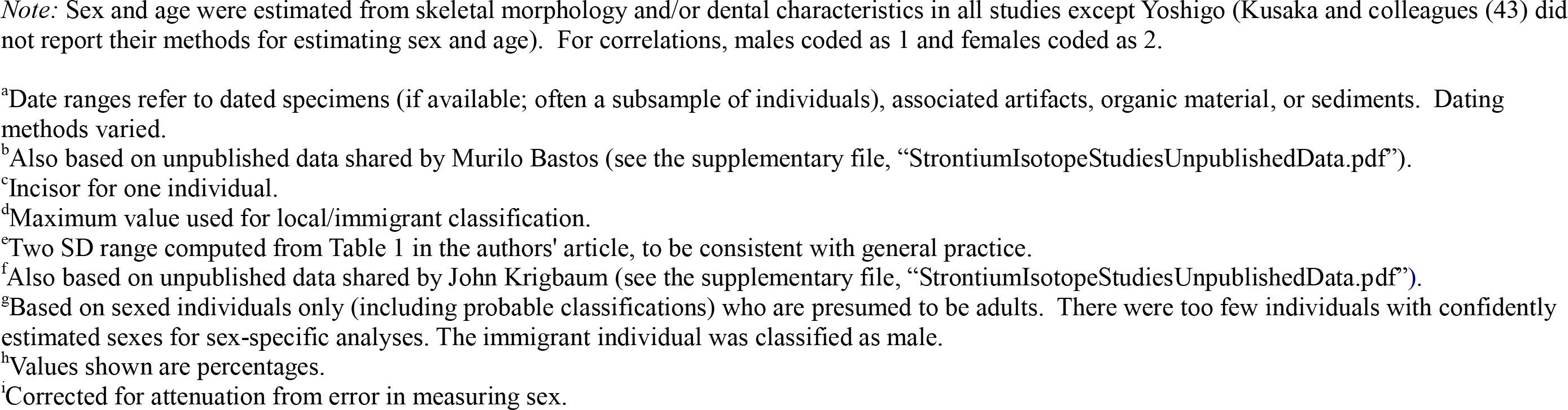

**Figure 2.**
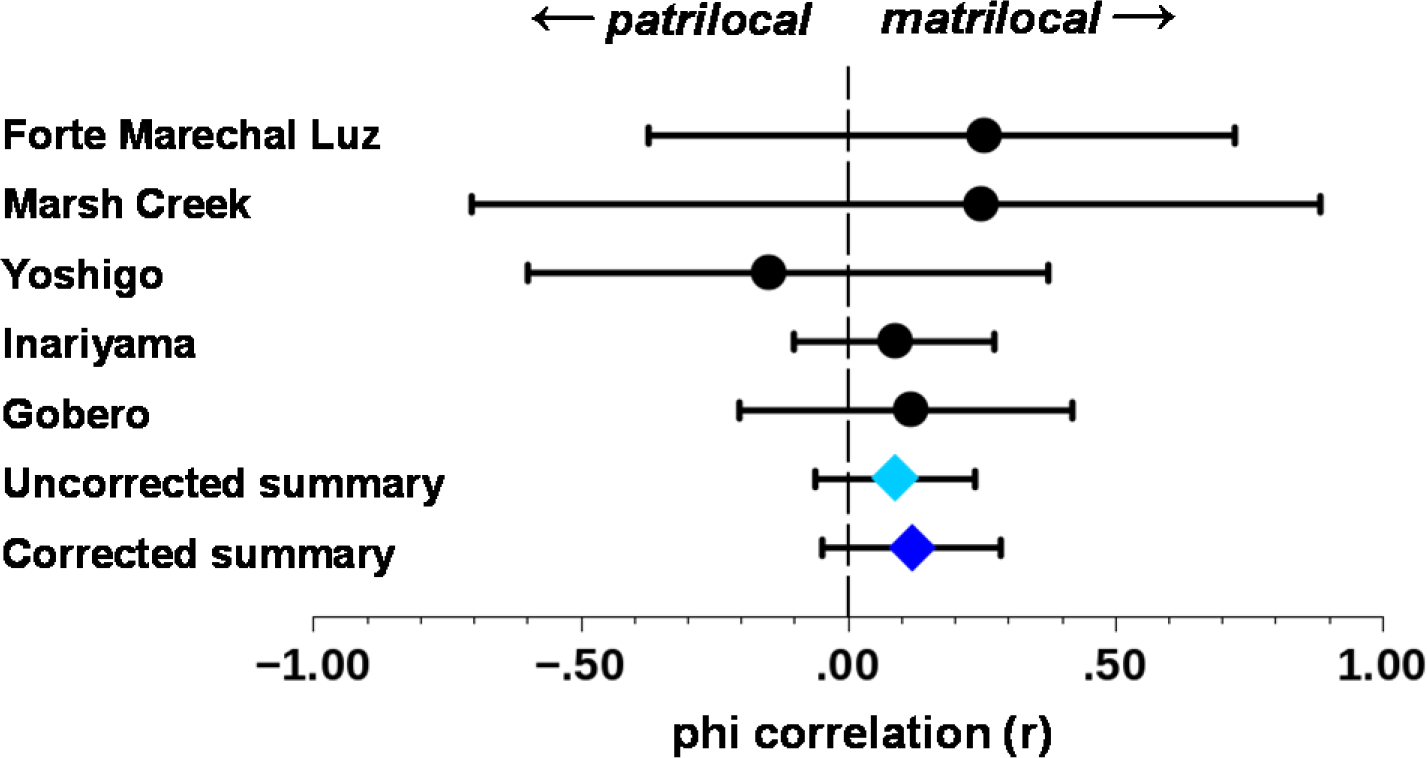
Associations between sex and local/immigrant status, strontium isotope studies. Bars show 95% confidence intervals.

**Table 2.** Association between sex and deviation from the local strontium isotope range in immigrant individuals

| Site | *n* immigrants (*n* male/*n* female) | *r* (95% confidence interval) |
| --- | --- | --- |
| Gobero, Niger (19,20) | 5 (3/2) | .32 (-.93 - .98) |
| Inariyama, Aichi Prefecture, Japan (22) | 4 (2/2) | .77 (-.74 - .99) |
| Marsh Creek, San Joaquin River Delta, California, USA (23) | 35 (22/13) | -.16 (-.47 - .18) |
| Yoshigo, Aichi Prefecture, Japan (25) | 14 (9/5) | -.01 (-.54 - .52) |
| *Summary* |  | -.09 (-.36 - .20) |
| *Summary corrected for attenuation<sup>a</sup>* |  | -.12 (-.47 - .26) |

*Note:* Males coded as 1, females coded as 2.
<sup>a</sup>Corrected for attenuation from error in measuring sex.

#### Other modern human studies

Other studies met the inclusion criteria, but could not be included in the meta-analyses because large proportions of individuals showed mobility during childhood, hindering a clear interpretation residence patterns. Haverkort and colleagues (44,45) studied strontium isotope ratios in 54 individuals buried at three different sites, dated to 8000-4000 years B.P., near Lake Baikal in Siberia. They measured these ratios in the enamel of first, second, and third molars as well as in femur bone samples. Haverkort and colleagues did not definitively classify individuals as locals or immigrants, and consequently, it is not possible to estimate post-marital residence patterns from their data. The local isotope ratios in terrestrial and aquatic animals differed substantially both within and between geologically defined areas, hindering the definition of narrow local isotopic ratio ranges. Isotope ratios varied dramatically within individuals across different molars and adult femur bone. Weber and Goriunova (46) did classify individuals (*n* = 19 adults) from one of these sites as locals or immigrants based strictly on first molar (infancy) strontium isotope ratios. This classification, however, is problematic for post-marital residence patterns. Fifteen (79%) of the 19 adults had at least one non-local isotope ratio at some point in childhood/early adolescence. Half (3/6) of the juveniles also had at least one non-local value. About half of enamel values were in the local range in infancy (5/9 men, 0/1 women), early childhood (5/9 men, 1/1 women), and late childhood/early adolescence (3/9 women, 0/1 women). Five (42%) of the 12 individuals with a local value in infancy had a subsequent non-local value in childhood/early adolescence. Furthermore, all individuals, including juveniles, had isotopic ratios for bone outside of the local range defined by modern fauna and flora. Weber and Goriunova noted that varying seasonal and inter-annual migration patterns between geochemical regions could account for the inconsistent ratios within and between individuals.

Harald and colleagues (47) conducted a similar investigation with results that are also difficult to interpret for post-marital residence patterns. They studied post-marital residence patterns in 50 individuals buried at the Cecil (Bear Creek; CA-SJO-112) site near Stockton, California, USA, which was dated to 3260-2950 years B.P. (uncalibrated). Harald and colleagues measured strontium isotope ratios in enamel in first molars/canines/incisors, second molars, and third molars, as well as in cortical bone. They estimated the local range of isotopic ratios from the mean ratio in bone ± 2 standard deviations. Harald and colleagues noted the similarity between this range (0.70656-0.70677) and the 2 standard deviation range observed in the waters of the San Joaquin Delta about 20-30 km downstream (0.70574-0.70718) (48). Of the 34 adults with isotope ratios measured for enamel, 26 (76%) had at least one non-local value during childhood and/or adolescence. Men were more likely to have local values in infancy (first molars/canines/incisors) (9/11) than women (2/9). From early childhood to adolescence, very few men or women had local values (early childhood/second molars:1/6 men, 0/5 women; late childhood/adolescence/third molars:1/9 men, 1/5 women). Of the 13 adults who had local values in infancy and another isotope measurement from molar enamel, most (6/8 men, 1/1 woman, 1/4 sex unknown) had a non-local value later in childhood and/or adolescence.Incidentally, all of the isotope ratios in enamel and bone that Harald and colleagues measured fell within the local range for the San Joaquin Delta waters. Moreover, this site is only about 36 kilometers (geodesic distance) from a concurrent site, Marsh Creek, which appears to have been moderately endogamous with matrilocal tendencies (Table 1).

Harald and colleagues interpreted their results as indicating patrilocality, with bride service being performed by most males starting in early childhood. To my knowledge, there is no ethnographic precedent of bride service beginning before adolescence. Regardless, it seems that raising a young child—a potential future son-in-law--would be more of a burden to a woman’s parents than a service. Another post hoc interpretation parallels that of Weber and Goriunova (46) for their similar profile of results. The strontium isotope ratios in bone could have been affected by diagenesis, whereby the bone was contaminated with strontium from the soil and groundwater at the site after burial, producing an artifactually narrow band of ratios (29). The variation in individuals’ isotope ratios in childhood and adolescence might then reflect movement by a single community within a larger territory, perhaps to seasonal or multi-annual camps. Multi-annual camps or subsets of the community alternating between camps over time, in particular, might account for much of the variability between individuals.

Eerkens and colleagues (49) studied post-marital residence patterns in 14 adults buried at the Sand Hill Road site (CA-SCL-287) near Stanford, California, USA, which was dated to 1987-1301 years B.P. Their results are almost a mirror image of those reported by Harald and colleagues (47) at the Cecil site, approximately 105 kilometers away (geodesic distance, www.sunearthtools.com. Eerkens and colleagues measured strontium isotope ratios in enamel in first molars (infancy) and third molars (late childhood/early adolescence), as well as in cortical bone. Of the 13 sexed adults, 7 (54%) had at least one non-local value during childhood or early adolescence, based on a ±2 standard deviation local range of the isotopic ratios in bone (from my calculations with data reported in their Table 1, given discrepancies in the authors’ classifications). Women (6/6) were more likely to have local values in infancy than men (3/7). During late childhood/early adolescence, most adults had non-local values (6/6 men, 1/3 women). Half of those with a local value in infancy and an isotope measurement for their third molars had a non-local value during late childhood/early adolescence (1/3 women, 2/3 men). Eerkens and colleagues interpreted these results as indicating matrilocality. However, as with the Cecil site, the local range of isotopic ratios could be artifactually restricted by diagenetic contamination of the cortical bone. Consequently, much of the variation in isotopic ratios in enamel between and within individuals might reflect normal movement within the community’s territory.

#### Extinct hominins

In 2011, Copeland and colleagues (50) reported their isotope analyses of fossil canine and molar enamel from 19 hominin (Australopithecus africanus and Paranthropus robustus) individuals found in the Swartkrans and Sterkfontein caves in South Africa. The individuals were dated from 2.2 to 1.8 million years B.P. Based on the moderate sexual dimorphism in modern primate tooth size, they inferred that individuals with larger teeth were males, and those with smaller teeth were females. They also implicitly assumed all specimens represent adult individuals, apparently not allowing for the possibility that any teeth came from juveniles. Copeland and colleagues classified 5 of 10 smaller tooth individuals, but only 1 of 9 larger tooth individuals, as immigrants (r = -.42, 95% CI:-.73-.05). They interpreted these results as indicating patrilocality.

In addition to the unreliable method of determining the sex and age of the hominins (especially for canine teeth (51)), several other methodological matters render Copeland and colleagues’ interpretation of residence patterns premature. To construct the reference range for distinguishing local from immigrant individuals, Copeland and colleagues used isotope ratios from grasses found in a geologically-based swath that comprised about 60% of the area within 10 kilometers of the sites.hese grass ratios showed a fairly restricted range of values (0.721-0.734). Grasses from other areas only 2-3 kilometers from the sites, but not in this swath, had much different isotope ratios (range = 0.724-0.758). Thus, Copeland and colleagues assumed the individuals they analyzed had much smaller home ranges than, as they noted (50), extant primates that live in environments similar those in which the extinct hominins lived.

Price and colleagues (52) emphasized that in strontium isotope studies, “the use of prehistoric and/or modern samples of small animals is advocated in order to establish the biologically available level and to distinguish migrant individuals. Where possible, we recommend measurement of tooth enamel in fossil animals from the archaeological site under consideration” (p. 132). As other isotope studies of prehistoric hunter-gatherers in this review indicate, this approach is a common way to establish a valid local reference range. In a 2010 report, Copeland and colleagues (53) analyzed fossil rodent teeth, likely originally deposited by owls, from the same sites as the hominin specimens. They found a range of isotope ratios (0.725-0.789) that encompasses and extends well beyond the range observed for all of the hominins (0.727-0.745), despite the owls and rodents’ small home ranges (53). Incidentally, in their 2011 supplementary material, Copeland and colleagues (50) also reported isotope ratios for fossil tooth enamel from non-migratory antelopes with very small home ranges (54), baboons, rock hyraxes, and other mammals of roughly similar age from the same caves as the hominins. The ranges of isotope ratios for the antelope, baboon, and rock hyrax samples are 0.727-.745 (the same as for the hominins),0. 727-0.736, and 0.722-0.738, respectively. Therefore, it seems there is no solid empirical basis to classify any of the hominin individuals in their samples as immigrants to the local area around the caves.

Sillen and colleagues (55,56) analyzed the strontium isotope ratios in bone (cranial, mandible, and palate) fragments from 6 adult Australopithecus robustus individuals (1 male, 1 probable female, and 4 of unknown sex) found in the Swartkrans cave in South Africa. The only individual whose ratio (0.723) suggested non-local origin (even according to the local ratio ranges from archaeological and modern mammal enamel reported in Copeland and colleagues’ (50) supplementary material) was the male. As I noted earlier, bone samples are less reliable indicators of origin as bone remodels over time, while molar enamel is fixed in childhood and does not remodel (29). Altogether, the evidence from South Africa indicates a strong degree of endogamy and no detectable tendency toward patrilocality or matrilocality in extinct hominins.

## Reliability of Sex Classification Studies

### Methods

As I conducted the literature search on prehistoric hunter-gatherers’ post-marital residence patterns and read the reports, I learned that researchers’ estimations of sex based on bone morphology and dental characteristics could be uncertain and possibly unreliable. Because individual sex is a key variable in my overall project, I sought to assess the reliability of these estimates and adjust my summary of the strontium isotope studies accordingly.

#### Literature search

In January and February of 2016, I conducted a separate literature search for reports on the reliability of estimating the sex of prehistoric hunter-gather individuals. I focused on comparisons between estimates from bone and tooth morphology and those from amelogenin sequencing (a genetic measure of sex). I searched Scopus and Google Scholar with the following keywords:(gatherer OR collector OR forager OR paleo* OR archaic* OR mesolithic) AND (amelogenin AND morpholog*). As before,I also performed backward and forward citation searches (the latter with Google Scholar).

#### Inclusion criteria

I included reports if they had data on adult prehistoric hunter-gathers’ sexes, as estimated with two or more independent methods.

#### Data analysis

From each report, I included in my analyses individuals whose sex the authors classified confidently on morphologic criteria and who had amelogenin or other genetic sex classifications the authors considered to be sound. Initially I intended to compute the phi (Pearson) correlation between the two methods of classification for each report and produce a meta-analytic summary. As I collected and abstracted reports, I realized this approach was not viable. Some reports included too few individuals with which to calculate a correlation, and some reports included individuals from multiple sites. Consequently, it was not possible or meaningful to use the report as the unit of aggregation for my summary. Instead, I pooled observations across reports into a single cross-tabulation (morphologic vs. genetic methods) of individuals’ estimated sexes. The resulting phi correlation served as my reliability estimate. The difference between this correlation and 1 indicates the extent of measurement error.

I used this reliability estimate to correct for attenuation in the observed correlations between sex and local/immigrant status in each of the strontium isotope studies, using Spearman’s (57) formula. Then I performed a random effects meta-analysis on the disattenuated correlations, following the procedures I used for the unadjusted strontium results while incorporating Schmidt and colleagues’ (58) methods for summarizing corrected correlations. The disattenuation analyses do not account for any error in measuring individuals’ local or immigrant statuses (error in measuring strontium isotope ratios or defining local ratio ranges).

### Results

#### Literature search and study characteristics

Figure 3 shows the results from my literature search on the reliability of sex classification. Records identified in the initial search were unduplicated within, but not between databases, and records from forward citation searches were unduplicated within, but not between, specific searches. A supplementary file (ReportsExcludedFromMetaAnalysisOfMeasurementOfSex.pdf) for this article lists the citations to retrieved reports I excluded from the meta-analysis and my reasons for excluding them.

I found 7 reports that included both morphologic and genetic assessments of sex (Figure 3 and Table 3). These reports included a total of 61 individuals from sites on every inhabited continent except Africa and Australia. The dates of these sites are comparable to those in the strontium studies (generally several thousand years before present). The authors of these reports did not indicate how many raters or whether different raters made the morphological assessments. None of these studies were preregistered. Except for Gotherstrom and colleagues (59), all study teams received funding support from national government agencies and most also had support from universities and/or private foundations.

**Figure 3.**
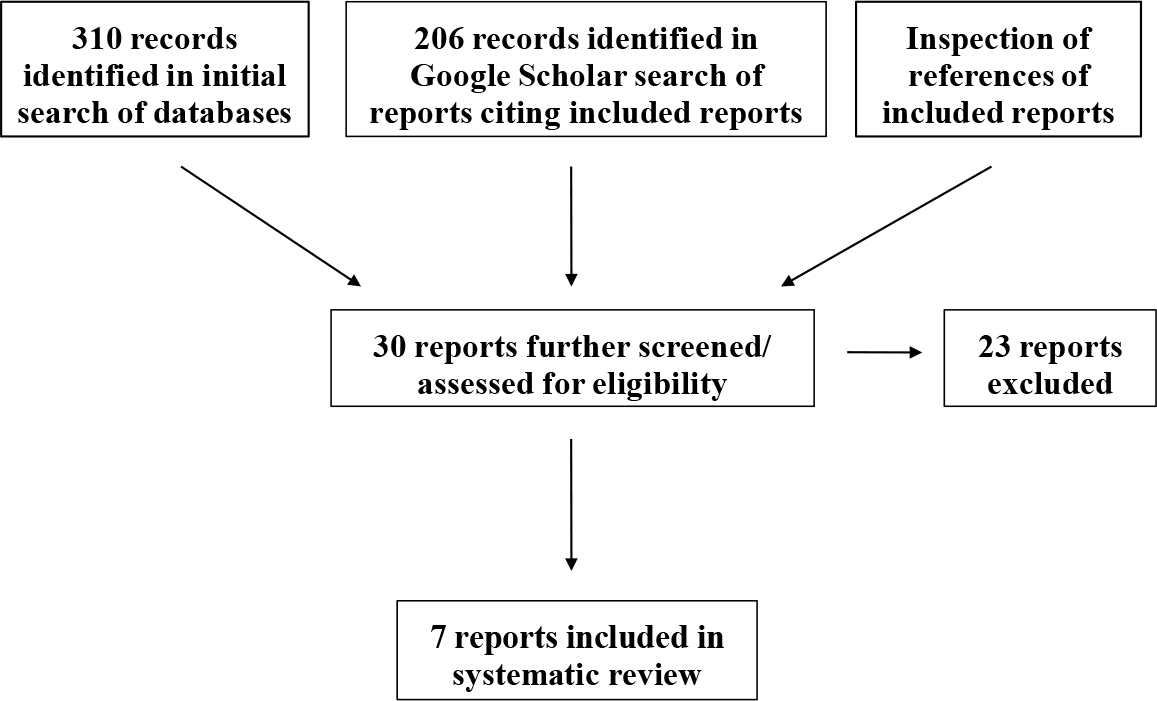
Results of the literature search for reliability of sex classification studies

**Table 3.**
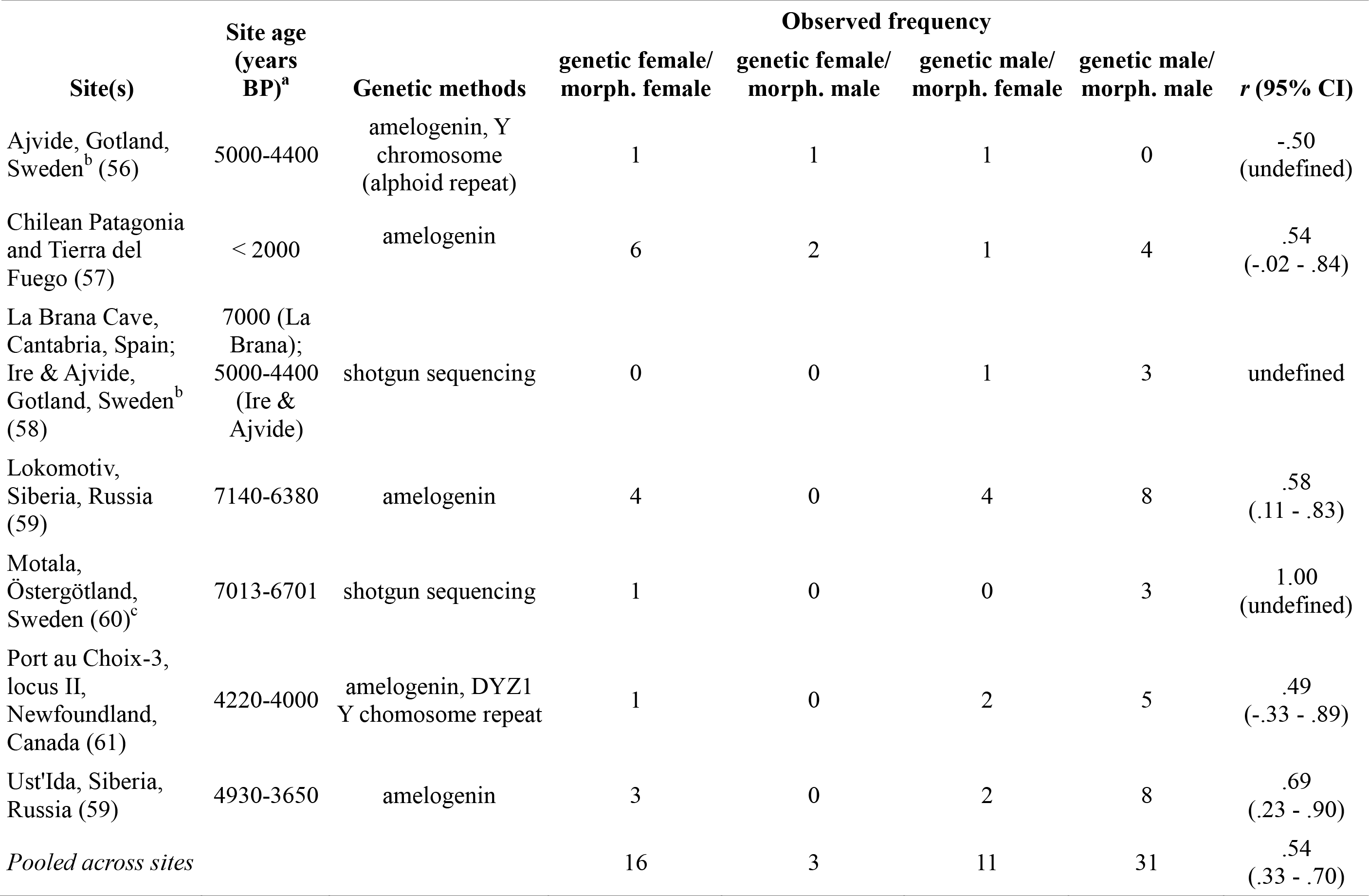
Reliability of measuring sex in prehistoric hunter-gatherer individuals:genetic vs. morphologic methods

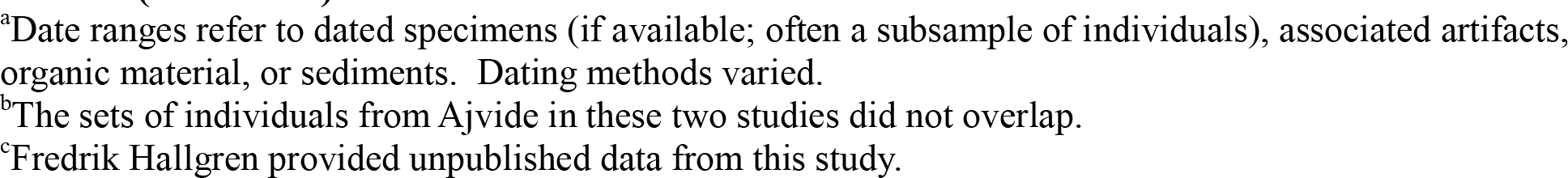

#### Reliability estimate

The pooled correlation between morphologic and genetic methods of sex estimation is.54 (95% CI:.33−.70). Nearly one-quarter of raters’ morphology-based sex estimations were inconsistent with the genetic measurements. Individuals assessed morphologically as male were quite likely to register as male by the genetic assessment (positive predictive value = 31/34 =.91), while a relatively modest proportion of individuals assessed morphologically as female registered as female by genetic assessment (positive predictive value = 16/27 =.59).

#### Correcting strontium correlations for attenuation from measurement error

All strontium isotope studies included in my review involved morphologic criteria for estimating sex, when authors mentioned a method in their reports. After correcting for attenuation from error in measuring sex, the matrilocal tendency in the strontium isotope studies increased slightly (corrected mean phi correlation =.12, 95% CI:-.05-.28; Tables 1 and 2).

## Mitochondrial DNA Studies

### Methods

mtDNA is maternally inherited in humans. mtDNA is often recoverable from archaeological bone, tooth, and other preserved tissues, even after many thousands of years, depending on the events and processes affecting the remains after death. A person’s mtDNA can be classified at both general and specific levels. mtDNA haplogroups are broad classifications, representing major branches of the mitochondrial phylogenetic tree. Individuals with the same haplogroup may have matrilines that join within as many as hundreds or perhaps even thousands of prior generations. mtDNA haplotypes, in contrast, are very specific classifications, representing the leaves or nodes near the leaves of the mitochondrial phylogenetic tree. Individuals with the same mtDNA haplotype likely have matrilines that join within only a few prior generations. Some aspects of postÒmarital residence patterns can be inferred from comparing mtDNA distributions between women and men, both within and between communities. mtDNA distributions are also not affected by whether the burial community was sedentary or nomadic.

Researchers typically measure mtDNA haplotypes by sequencing a hypervariable region of the mitochondrial genome that is prone to mutation. The high rate of mutations in this region means that occasionally individuals with matrilines that join within a few prior generations (or even one prior generation-siblings or mother and offspring) have different haplotypes (65,66). The high mutation rate also occurs within individuals due to unrepaired or incorrectly repaired DNA damage and during DNA replication in cell division over the life span, producing diversity in mtDNA haplotypes within individuals (67,68). Consequently, individuals with similar, yet distinct, haplotypes still may have closely related matrilines.

Measuring the similarity of haplotypes, or the inverse of genetic distances between them, lessens somewhat the problems of relying on qualitative classifications of haplogroups and haplotypes. Individuals with relatively close maternal kin connections, even if they do not share the same haplotype, have very similar haplotypes (corresponding to small genetic distances among them). However, variation in larger genetic distances (e.g., similarities between haplotypes in different haplogroups) often may not be meaningful in assessing post-marital residence patterns as such distances represent differences in degree of very remote matrilineal ancestry. Because haplogroups, haplotypes, and genetic distance each have strengths and weaknesses as levels of measurement, I focus on all three, without prioritizing one over the others.

#### Literature search

The general searches I used for the strontium isotope studies also applied to the mtDNA studies.

#### Inclusion criteria

I included an mtDNA report in the review if the sample had at least four adult (estimated age > 15 years) individuals, including at least two males and two females (based on confident, not probable, sex estimations). For one particular kind of analysis, I included contemporaneous neighboring sites if each site had the necessary data on at least two adult females. I included individuals in analysis if their haplogroups/haplotypes were confirmed by at least double extraction.

#### Data analysis

Measures of mtDNA diversity within a community, whether overall or by sex, are not useful for assessing post-marital residence patterns. Low diversity does not necessarily indicate matrilocality. If neighboring communities that exchange mates had similar mtDNA distributions, perhaps resulting from common ancestors for both communities, even patrilocality would still produce low mtDNA diversity within communities. Similarly, high diversity does not indicate patrilocality, as a community could have practiced endogamy or matrilocality yet also have descended from an ancestral population with high mtDNA diversity.

Another critical fact is that the mtDNA distribution of native members of a community is determined by the mtDNA distribution of mothers in that community. For instance, with patrilocality, men’s mtDNA distribution lags women’s distribution by a generation, on average.

I developed a few rules for defining haplogroups and haplotypes when reports were not entirely clear.If authors classified some individuals into an “other” haplogroup category and haplogroup-defining mutations in these individuals were not identical, then I treated each individual in the “other” category as belonging to a unique haplogroup (different from all others in the “other” category and other individuals in the sample). I used authors’ definitions of haplotypes when they defined them.Otherwise, I used reported data on individuals’ mtDNA variants at particular nucleotide positions along with their haplogroup designations to determine whether pairs of individuals had identical sequences for those regions and positions sequenced. In such cases, I considered this most specific level of sequence information to represent haplotypes. Moreover, I analyzed individuals’ haplotypes and haplotype similarity only when haplotype data were available for all individuals for whom haplogroups were ascertained.

I compared women’s and men’s mtDNA distributions both within and between burial sites. Each kind of comparison provides somewhat different evidence about post-marital residence patterns. Both kinds of comparisons involve computing a correlation between a hypothesis or structure matrix and a data matrix. These square matrices indicate hypothesized and observed similarity, respectively, of mtDNA classifications for pairs of individuals.

*Within-site comparisons*. For each site, I constructed up to three data matrices that represent the matrilineal similarity between individuals, depending on the available data. For haplogroups and haplotypes, the data matrices are binary (values of 1 for pairs with the same haplogroup/haplotype, 0 otherwise). For haplotype similarity (inverse genetic distance), I created a valued data matrix representing the number of nucleotides each pair of individuals had in common, when assessed for positions in the mtDNA sequence where the sampled individuals exhibited variation. Figure 4 shows a hypothetical data matrix for a set of women and men found at a site. Each row/column represents a different individual. The rows and columns have been sorted by sex. There are four women and four men, and two of each sex have haplogroup A and two of each sex have haplogroup B.

**Figure 4.**
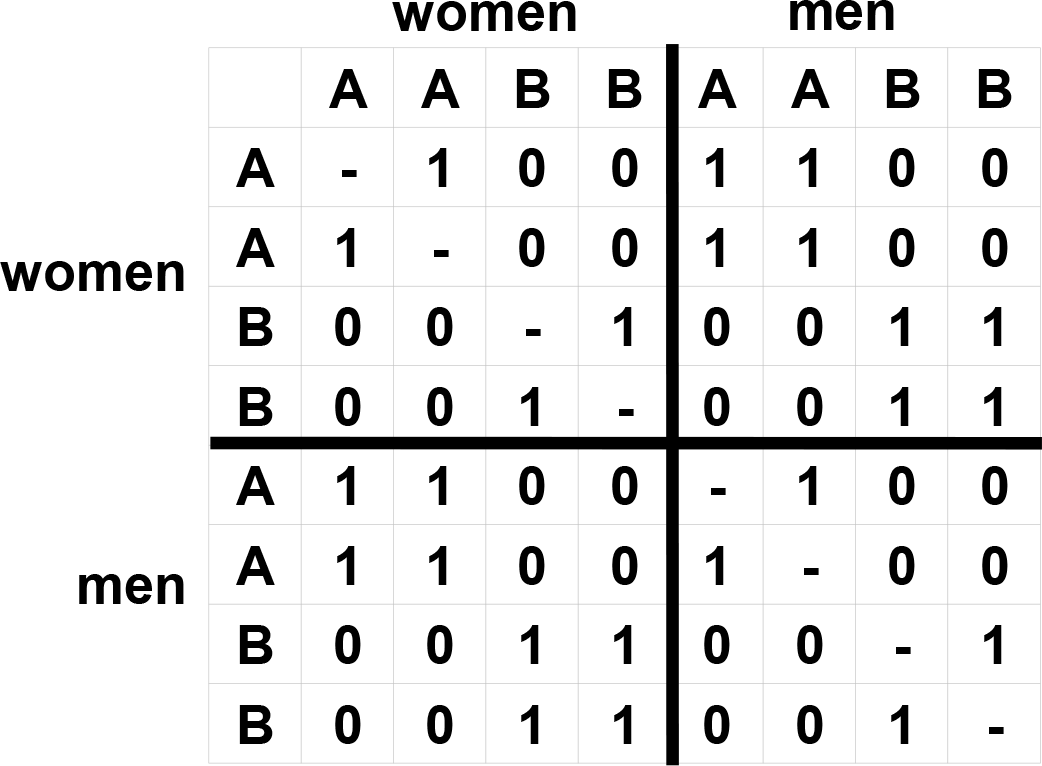
Hypothetical mtDNA haplogroup data matrix. Letters represent haplogroups.

Within-site comparisons involve assessing two closely related hypotheses (structures) that together allow matrilocality to be distinguished from other post-marital residence patterns. If, after marriage, women stay in their natal communities but men move to other communities where their wives live, then women should have more similar mtDNA to each other than they do with men or than men have with each other (as men might be drawn from multiple other communities). With matrilocality, then, women form a genetic core in the community and are more “central” if genetic similarities were conceived as a network. In Figure 5, a hypothetical structure matrix illustrates the idealized centrality pattern in which all the women have the same haplogroup/haplotype and each of the men has a haplogroup/haplotype different from all other individuals.

This structure matrix represents an extreme that is unlikely to be observed in real data. Under matrilocality, men’s mtDNA distribution would, over time, be a proportionate mixture of haplogroups/haplotypes of the women in the marriage-linked communities. Thus, men in a matrilocal community often would have some haplogroups/haplotypes in common with each other and with women, but at a lower rate than women have with each other. Figure 5 reflects that expectation.

**Figure 5.**
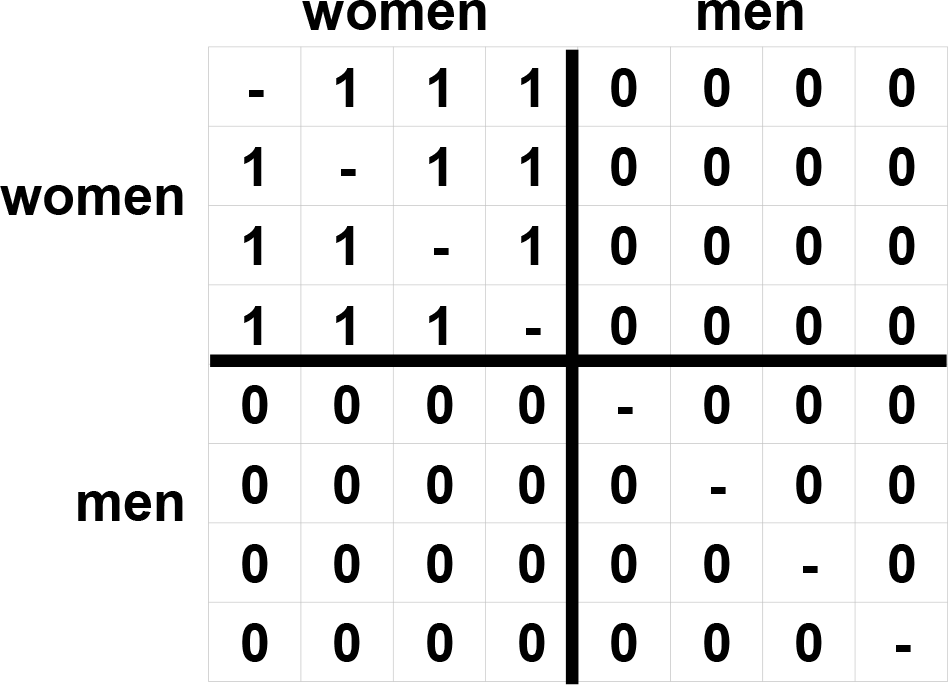
Hypothetical centrality hypothesis structure matrix (within-site comparisons)

To measure the correspondence between a structure matrix and a data matrix, I computed a matrix correlation (Pearson), with each matrix treated as a vector. For the centrality hypothesis, a positive correlation indicates a matrilocal tendency. A negative correlation, however, is difficult to interpret in isolation. For instance, the data matrix in Figure 4 represents identical mtDNA distributions for women and men, reflecting no matrilocality. Yet the correlation between the centrality hypothesis and this data matrix is negative, because the main diagonal, which represents each individual’s similarity to self, is ignored. In this circumstance, cross-sex pairs have greater similarity on average than within-sex pairs.

Therefore, for each site I also constructed a second structure matrix that is the same as that for the centrality hypothesis except that the cross-sex pairs are ignored (as structural zeros) (see Figure 6).The correlation between this matrix and the data matrix (with cross-sex pairs excluded) represents a comparison just between the mitochondrial similarity between women with that between men. I call this the “density” correlation, as it refers to whether women have greater similarity (shorter genetic distances) with each other than men do with each other. Positive values indicate matrilocality, although negative values are not necessarily inconsistent with matrilocality.

**Figure 6.**
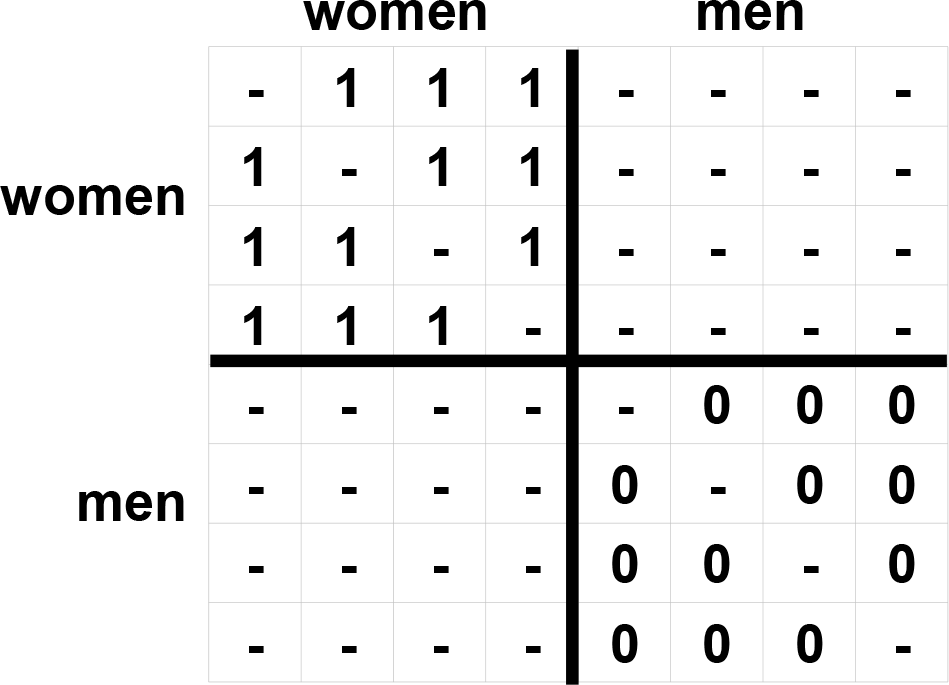
Hypothetical density hypothesis structure matrix (within-site comparisons)

Taken together, the centrality and density correlations imply post-marital residence patterns crudely. If either or both correlations are positive, it suggests matrilocality. If both correlations are negative, it suggests either endogamy, patrilocality, or bilocality. For the centrality and density comparisons, these latter three post-marital residence patterns are indistinguishable (this is so for patrilocality after a few generations).

For each matrix correlation I calculated, I also performed the Quadratic Assignment Procedure (QAP) (69), which involves randomly permuting the rows and columns of the data matrix and recomputing the matrix correlation. I computed 10,000 permutations for each correlation, and from these results obtained a standard deviation of the permutation distribution and, by extension, a 95% confidence interval for the observed correlation.

The observed distributions of haplogroups, haplotypes, genetic distances, and sexes for a site usually constrain the range of possible correlation values greatly in both negative and positive ends of the spectrum. In practice, with real data, maximally strong relationships can yield modest correlation coefficients.

I had initially planned to produce a graphic display of mitochondrial similarities for each site with haplotype similarity data with multidimensional scaling. I abandoned this exercise as it did not enhance my interpretation of the results beyond the analyses I have already described.

When authors reported both morphologic and genetic estimates of individuals’ sexes, I used the genetic estimates for defining sex in my main analyses. I also carried out separate within-site analyses for the different methods of sex estimation. I performed these analyses for just those individuals with sex estimations from both methods. I planned these additional analyses before conducting them, but I neglected to include them in an updated version of my project protocol.

*Between-site comparisons*. mtDNA studies of burial sites that were contemporaneous, geographically close, and similar in ecology and material culture provide additional evidence on post-marital residence patterns in prehistoric hunter-gatherers. Such sites likely represent communities that were in direct contact with each other. Over time, such contact might have included exchange of mates.

To make the samples from each community comparable temporally, I included all individuals from one site whose estimated dates (+/– 2 SD) fell between the earliest and latest dates (+/– 2 SD) for individuals from the other site(s), and vice versa. I assumed that individuals with no dates fell within the observed range of estimated dates for other individuals at the site. Also, if few individuals at a site had estimated dates, I used the estimated dates for other materials at the site as the reference range for that site. I considered inter-site distances of 150 km or less to be small enough to assume at least occasional contact between the burial communities.

I examined how dissimilar mtDNA distributions were between sites, with women and men analyzed separately (when both morphologic and genetic estimates of sex were available, I used genetic estimates). Figure 7 shows the structure matrix for this kind of comparison. The structure matrix represents the hypothesis that women (or men) from one site have a homogeneous mtDNA distribution that is entirely distinct from women (or men) from another site. The structure matrix can be expanded accordingly for comparisons involving more than two sites. For each set of sites, I calculated the matrix correlations between the corresponding structure and data matrices, as well as QAP tests with 10,000 permutations.

**Figure 7.**
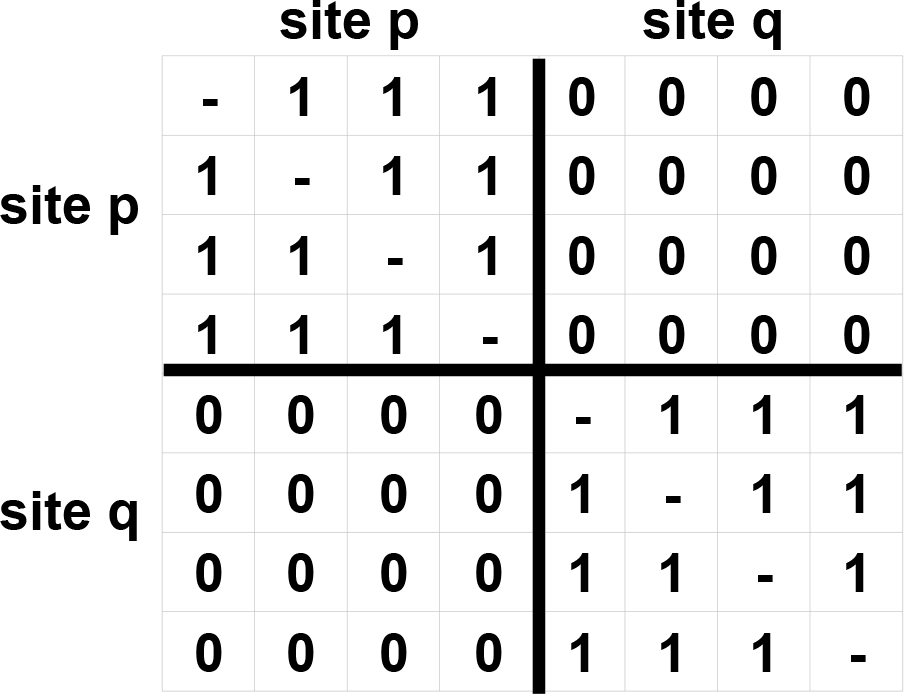
Hypothetical between-site structure matrix (for either women or men)

A negative inter-site correlation for women indicates some degree of female exogamy. Such female exogamy could have co-occurred with some or an even greater degree of male exogamy, but this comparison cannot reveal such patterns. A positive inter-site correlation for women and a negative correlation for men indicates matrilocality. Positive correlations for both women and men indicate endogamy. These interpretations are analogous to Aguiar and Neves’ (70) and Williams and colleagues’ (28) interpretations of parallel inter-community analyses.

*Meta-analysis*. I summarized the correlations for each type of comparison with random effects models and forest plots (31). The conventional approach for estimating the standard error of a correlation is based on the associated sample size. For matrix correlations, this would be inappropriate, because the unit of analysis is the pair of individuals. Therefore, I used the standard deviation of the QAP permutation distribution to estimate the standard error of an observed matrix correlation.

The within-site and between-site analyses each involved two comparisons (centrality and density hypotheses for within-site, separate correlations for women and men for between-site). To apprehend the results more easily, I displayed these series of paired correlations in scatterplots. I drew 95% confidence ellipses (based on the 95% confidence intervals for the correlations in each pair) around each data point in these graphs. These ellipses are crude, because they do not account for the dependence between the paired correlations.

### Results

#### Literature search and study characteristics

The literature search for the mtDNA studies was part of the general search that included the strontium isotope studies (Figure 1). Table 4 shows key details of the mtDNA studies of anatomically modern humans included in my main review. There are many sites from each of Asia, Europe, and North America. The sites date from several thousand to a few hundred years before present. Authors used genetic methods for estimating the sex of some individuals from Lokomotiv, Port au Choix-3, Shamanka II, and Ust’Ida. Authors estimated the sex of other individuals from these sites and all individuals from the other sites in Table 4 based on morphologic and dental characteristics only. Just three teams of authors considered post-marital residence patterns in their analyses or interpretations.

**Table 4.**
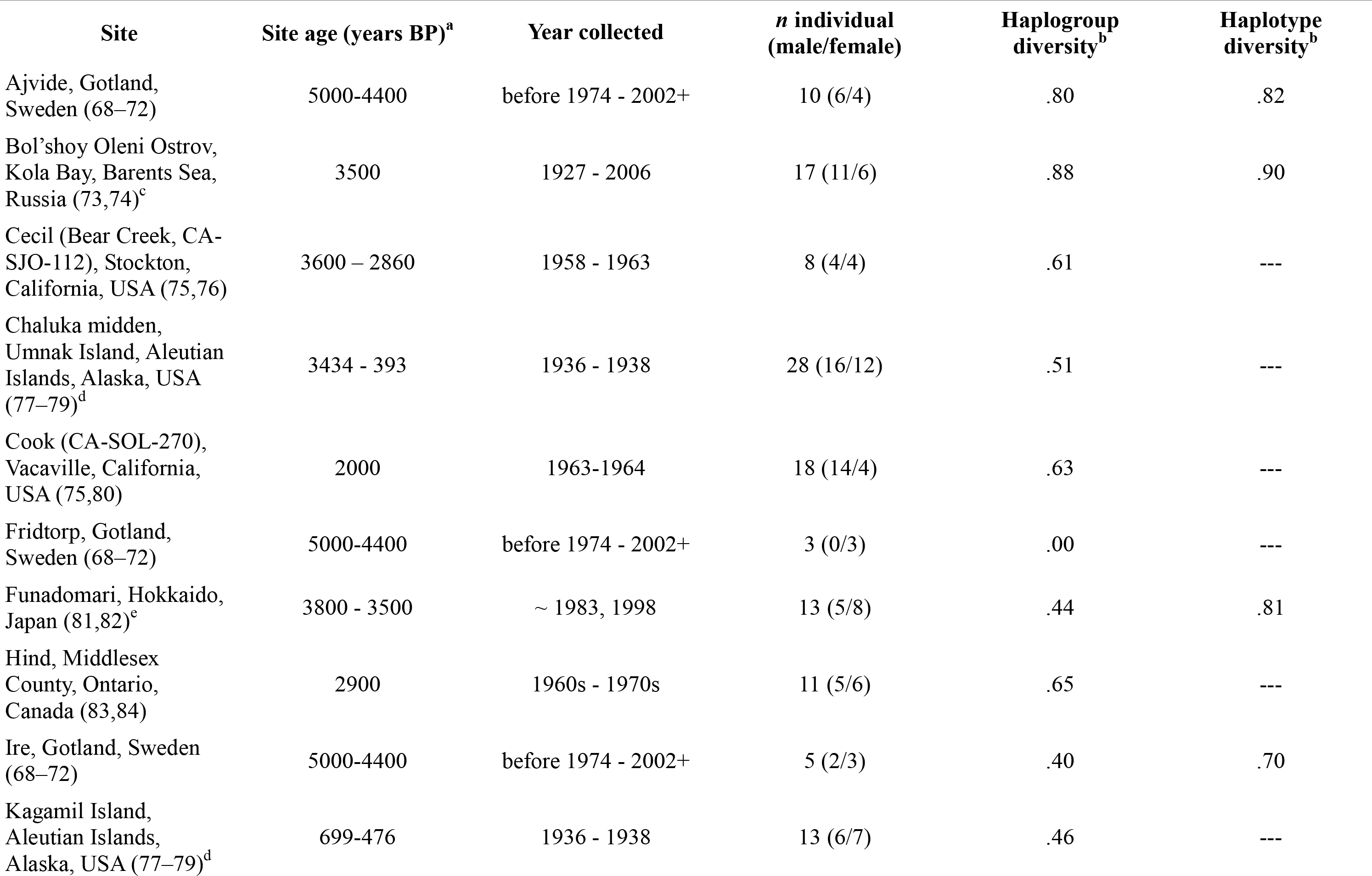
mtDNA studies

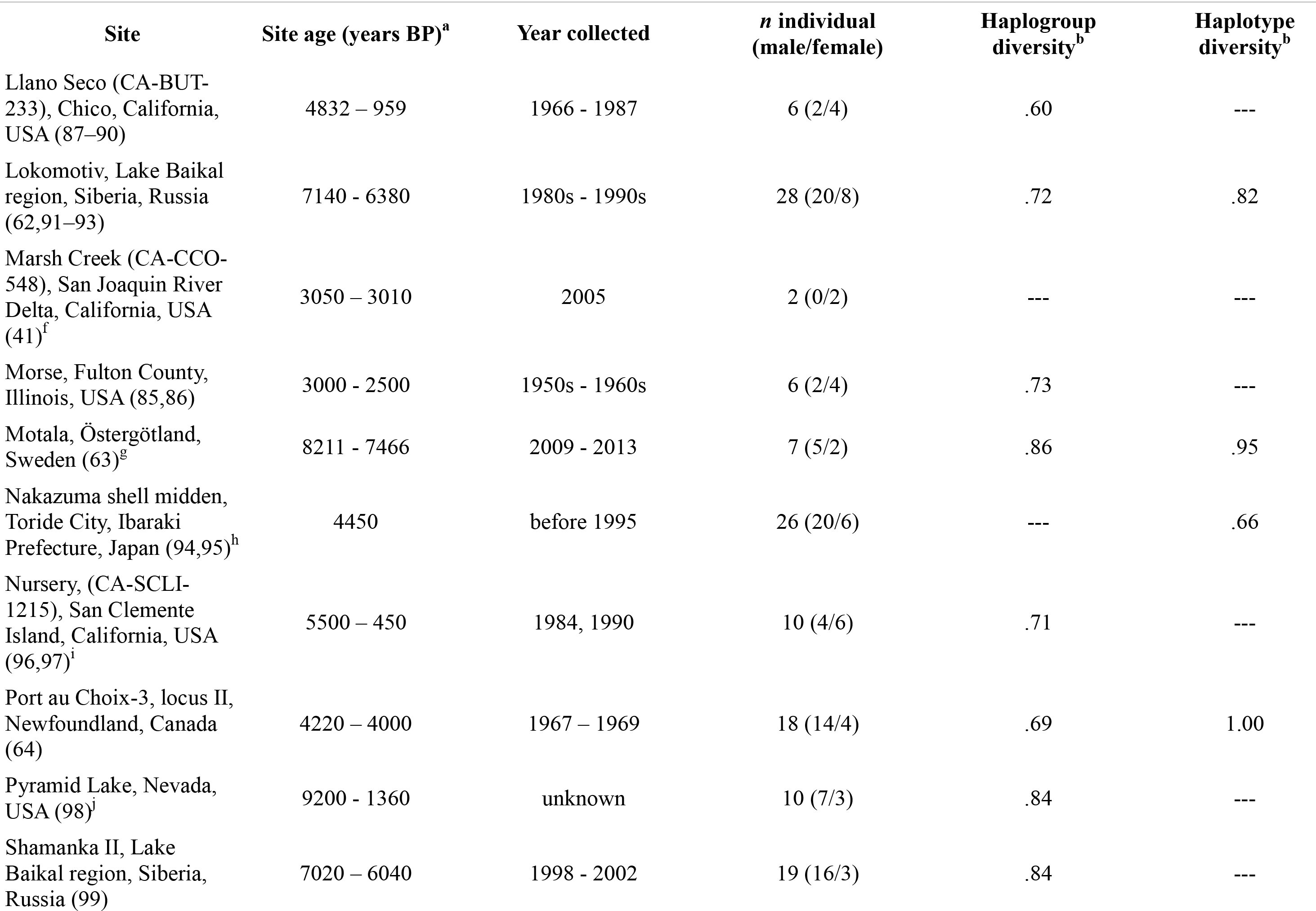

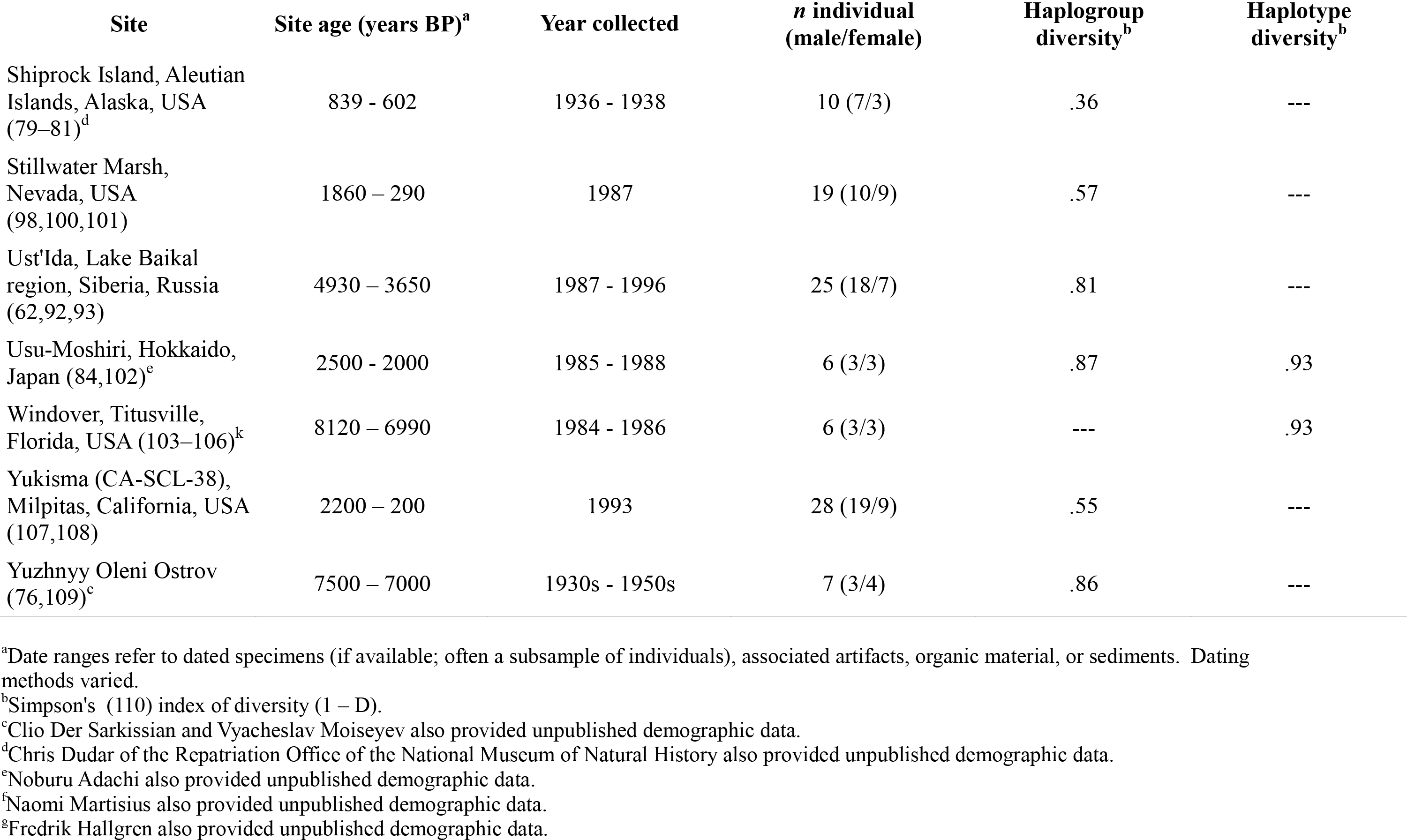

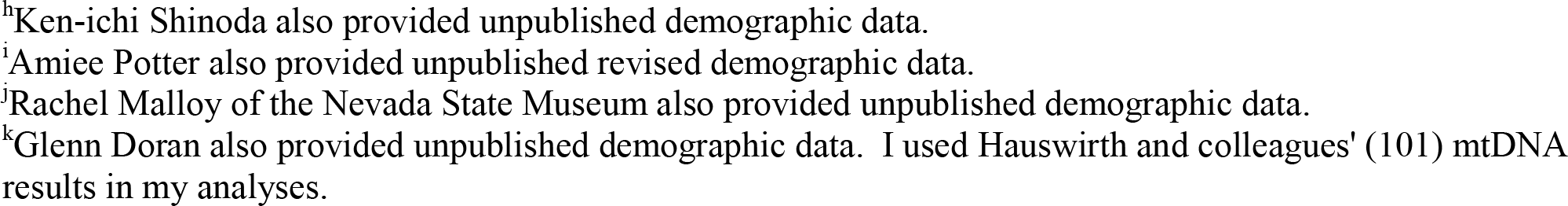

All unpublished data from the mtDNA studies included in my analyses are in a supplementary file (mtDNAStudiesData.xls). Another supplementary file (ReportsExcludedFromSystematicReviewOfResidencePatterns.pdf) for this article lists the citations to retrieved reports of mtDNA studies I excluded from the review and my reasons for excluding them.

None of the studies were pre-registered. Most authors received funding from government agencies to carry out their studies, and some had additional funds from private foundations or universities. Several authors did not report a funding source.

Nearly all authors reported attempts to prevent and detect contamination in their specimens. Naturally, the sophistication of these procedures varied over time, with earlier investigations usually somewhat less rigorous by current standards than more recent studies. Authors who assessed haplotypes sequenced HVR1 (hypervariable region 1) of the mitochondrial genome, although their precise definitions of this region varied somewhat in each study. Some authors also sequenced other regions or portions of other regions of the mitochondrial genome.

At most sites, there was moderate to high diversity in mtDNA haplogroups and/or haplotypes (Table 4). The diversity measure, Simpson’s (110) index of diversity (1-D), indicates the proportion of pairs of individuals with different haplogroups/haplotypes. The index ranges from 0 (all individuals with the same haplogroup/haplotype) to 1 (every individual has a different haplogroup/haplotype).

#### Within-site comparisons

Tables 5, 6, and 7 show the centrality and density hypothesis results at the level of haplogroups, haplotypes, and haplotype similarity, respectively. Figures 8, 9, and 10 show the corresponding results in graphic form. There was no genuine variation in the underlying summary estimates for the centrality hypothesis at any level and the density hypothesis at the haplogroup level (I^2^ =.00), but there was modest genuine variation for the density hypothesis at the haplotype and haplotype similarity levels (I^2^ =.13 and.35, respectively). The summary estimates of the centrality and density correlations at all levels hover around 0. These results indicate, then, no strong tendency toward any particular residence pattern.

**Table 5.** Within-site centrality and density matrix correlations for haplogroups.

| Site | N | Centrality r | Density r |
| --- | --- | --- | --- |
| Ajvide | 10 | -.03 (-0.31 - 0.25 ) | -.11 (-0.45 - 0.27) |
| Bol'shoy Oleni Ostrov | 17 | -.13 (-0.30 - 0.05) | -.20 (-0.45 - 0.07) |
| Cecil | 12 | -.06 (-0.37 - 0.27) | -.03 (-0.54 - 0.49) |
| Chaluka | 28 | .03 (-0.05 - 0.10) | .04 (-0.03 - 0.10) |
| Cook | 18 | -.08 (-0.24 - 0.08) | -.11 (-0.31 - 0.09) |
| Funadomari | 13 | -.07 (-0.35 - 0.24) | -.01 (-0.43 - 0.42) |
| Hind | 11 | -.10 (-0.40 - 0.21) | -.04 (-0.48 - 0.42) |
| Ire | 5 | -.36 (-0.84 - 0.45) | -.58 (-0.97 - 0.69) |
| Kagamil Island | 13 | -.13 (-0.40 - 0.15) | -.23 (-0.62 - 0.25) |
| Llano Seco | 6 | .17 (-0.10 - 0.41) | -.35 (-0.74 - 0.21) |
| Lokomotiv | 28 | -.02 (-0.16 - 0.13) | .01 (-0.21 - 0.22) |
| Morse | 6 | .43 (-0.02 - 0.74) | -.35 (-0.79 - 0.33) |
| Motala | 7 | -.09 (-0.49 - 0.33) | -.10 (-0.49 - 0.32) |
| Nursery | 10 | -.03 (-0.25 - 0.20) | -.22 (-0.49 - 0.08) |
| Port au Choix<sup>a</sup> | 18 | .08 (-0.14 - 0.29) | .13 (-0.20 - 0.43) |
| Pyramid Lake | 10 | .13 (-0.19 - 0.42) | .24 (-0.25 - 0.63) |
| Shamanka II | 19 | -.06 (-0.21 - 0.09) | -.07 (-0.25 - 0.10) |
| Shiprock Island | 10 | -.17 (-0.49 - 0.19) | -.27 (-0.75 - 0.39) |
| Stillwater Marsh | 19 | -.07 (-0.21 - 0.08) | -.08 (-0.30 - 0.14) |
| Ust'Ida | 25 | -.10 (-0.26 - 0.07) | -.15 (-0.38 - 0.11) |
| Yukisma | 28 | .00 (-0.19 - 0.18) | .00 (-0.31 - 0.31) |
| Yuzhnyy Oleni Ostrov | 7 | .04 (-0.45 - 0.52) | .25 (-0.51 - 0.79) |
| *Summary* |  | -.02 (-0.06 - 0.02) | -.02 (-0.07 - 0.03) |

*Note:* 95% confidence intervals of correlations are in parentheses. The centrality correlation for Usu-Moshiri is -.20 (95% CI: -.30 - .05; n = 6). No density correlation is calculable for this site because there were no same-sex pairs that had the same haplogroup. The total number of individuals in the studies summarized is 326.
<sup>a</sup>Jelsma (2000) did not classify three individuals into haplogroups; I treated each of these individuals as having a unique haplogroup.

**Table 6.** Within-site centrality and density matrix correlations for haplotypes

| Site | N | Centrality r | Density r |
| --- | --- | --- | --- |
| Ajvide | 10 | -.01 (-.31 - .29) | -.11 (-.52 - .35) |
| Bol'shoy Oleni Ostrov | 17 | -.12 (-.31 - .08) | -.17 (-.46 - .14) |
| Funadomari | 13 | -.03 (-.22 - .17) | .09 (-.17 - .35) |
| Ire | 5 | -.43 (-.85 - .32) | -1.00 (-.98 - .34) |
| Motala | 7 | -.05 (-.45 - .37) | -.10 (-.61 - .47) |
| Nakazuma | 26 | .15 (-.03 - .33) | .23 (-.05 - .47) |
| Windover | 6 | -.13 (-.58 - .37) | -.45 (-.88 - .38) |
| Yuzhnyy Oleni Ostrov | 7 | .04 (-.45 - .52) | .25 (-.51 - .79) |
| *Summary* |  | -.01 (-.11 - .09) | .01 (-.16 - .17) |

*Note:* 95% confidence intervals of correlations are in parentheses. The centrality correlation for Usu-Moshiri is -.13 (95% CI: -.58 - .37; n = 6). No density correlation is calculable for this site because there were no same-sex pairs that had the same haplogroup. No pairs of Port au Choix individuals had the same haplotype. The total number of individuals in the studies summarized is 91.

**Table 7.** Within-site centrality and density matrix correlations for haplotype similarity (inverse genetic distance)

| Site | N | Centrality r | Density r |
| --- | --- | --- | --- |
| Ajvide | 10 | .29 (.04 - .51) | .15 (-.16 - .44) |
| Bol'shoy Oleni Ostrov | 17 | .01 (-.27 - .29) | -.01 (-.46 - .45) |
| Funadomari | 13 | -.07 (-.37 - .24) | -.02 (-.46 - .43) |
| Ire | 5 | -.35 (-.83 - .45) | -.70 (-.98 - .56) |
| Port au Choix | 18 | .08 (-.10 - .26) | .10 (-.16 - .34) |
| Usu-Moshiri | 6 | -.13 (-.54 - .33) | .00 (-.54 - .54) |
| Windover | 6 | -.23 (-.68 - .34) | -.89 (-.98 - -.42) |
| Yuzhnyy Oleni Ostrov | 7 | -.10 (-.49 - .33) | -.06 (-.65 - .58) |
| *Summary* |  | .06 (-.06 - .17) | -.03 (-.25 - .19) |

*Note:* 95% confidence intervals of correlations are in parentheses. Haplotype similarity was not calculable for Motala or Nakazuma based on the information in their associated reports. The total number of individuals in the studies summarized is 82.

**Figure 8.**
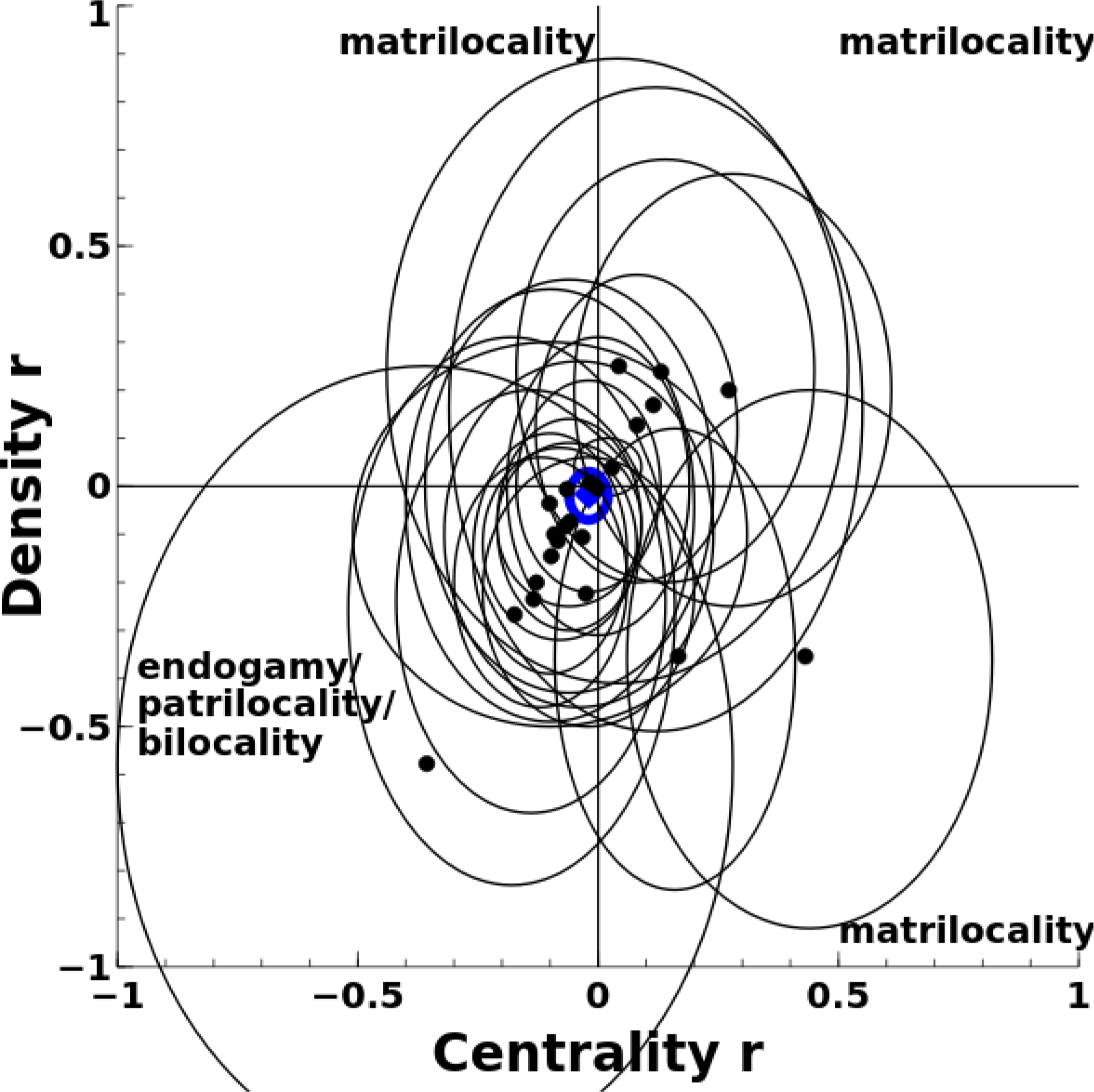
Scatterplot of centrality and density matrix correlations for haplogroups (within-site comparisons), including 95% confidence ellipses and summary estimate (blue, bold)

**Figure 9.**
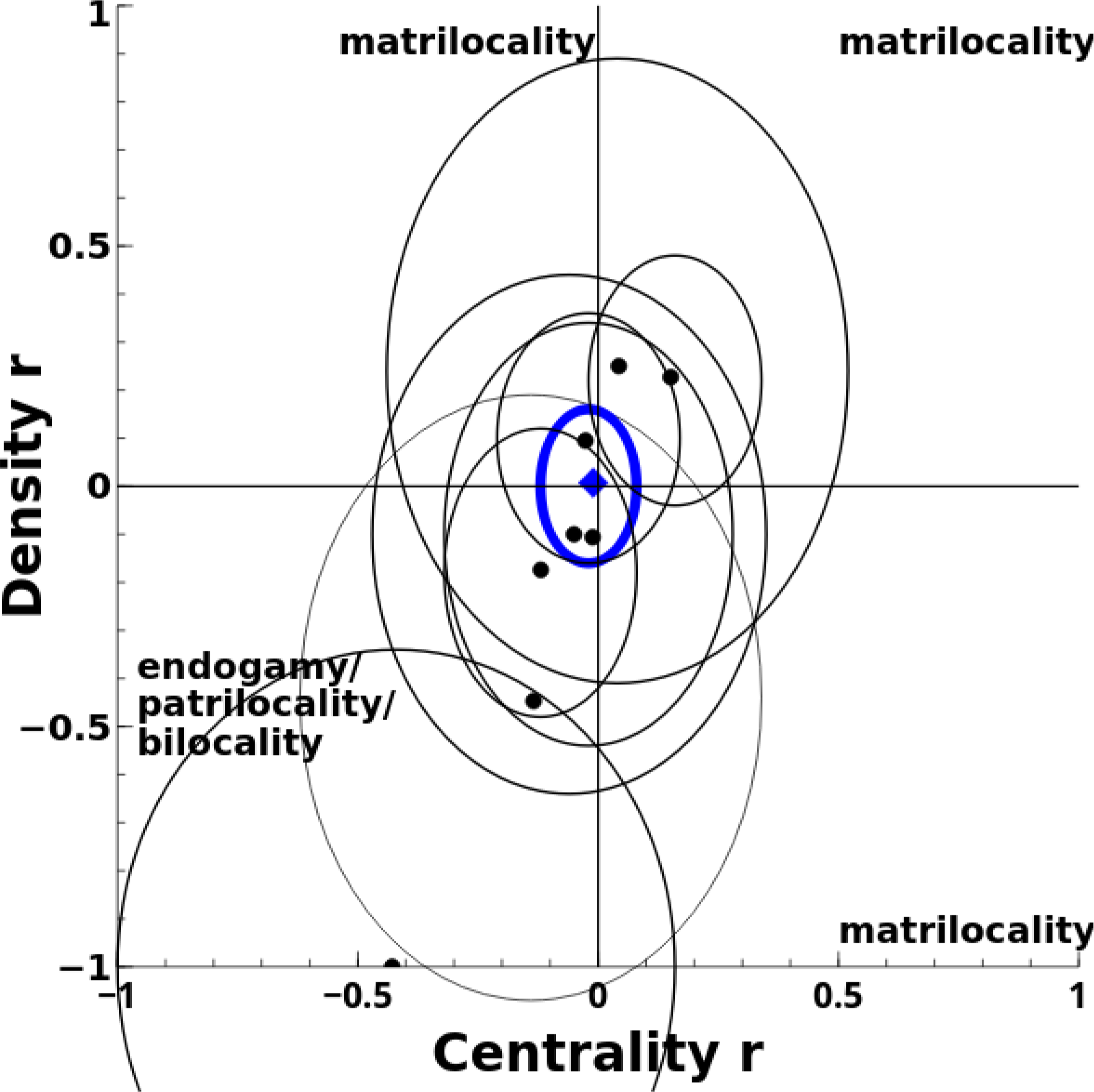
Scatterplot of centrality and density matrix correlations for haplotypes (within-site comparisons), including 95% confidence ellipses and summary estimate (blue, bold)

**Figure 10.**
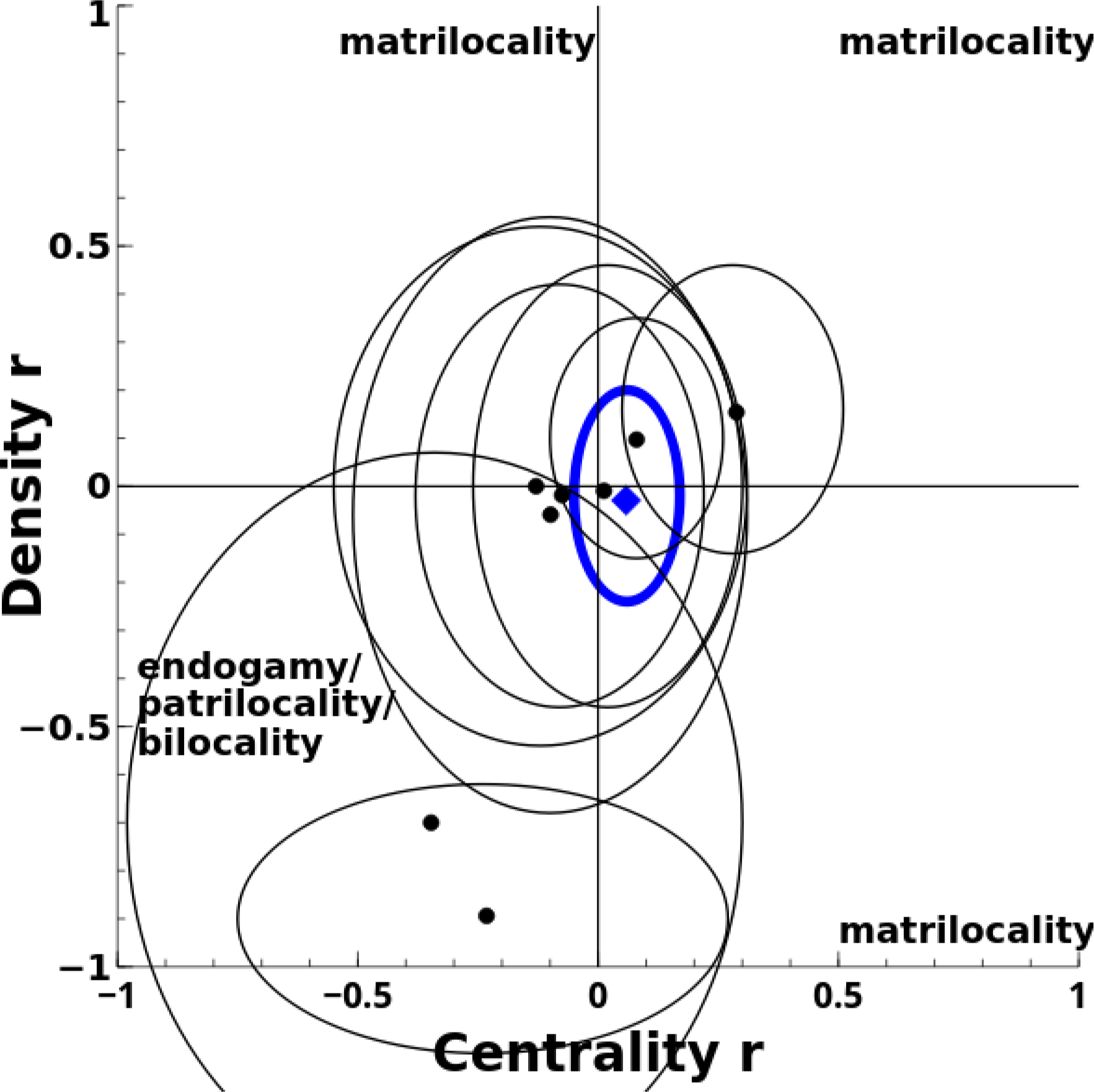
Scatterplot of centrality and density matrix correlations for haplotype similarity (within-site comparisons), including 95% confidence ellipses and summary estimate (blue, bold)

*Results by method of measuring sex*. Mooder (62) and Mooder and colleagues (91) estimated the sex of individuals from Lokomotiv and Ust’Ida with both morphologic and genetic methods. For both sites, morphologic estimates of sex produce results slightly in the direction of endogamy, patrilocality, or bilocality, while genetic estimates of sex produce results weakly in the direction of matrilocality (Figure 11). For Lokomotiv, at the haplogroup level, the within-site correlations based on morphologic and genetic sex estimations, respectively, are -.09 and.00 (for the centrality hypothesis) and -.08 and.03 (for the density hypothesis) (*n* = 16). The within-site correlations at the haplogroup level for Ust’Ida based on morphologic and genetic sex estimations, respectively, are −.06 and.10 (for the centrality hypothesis) and −.06 and.16 (for the density hypothesis) (*n* = 13).

**Figure 11.**
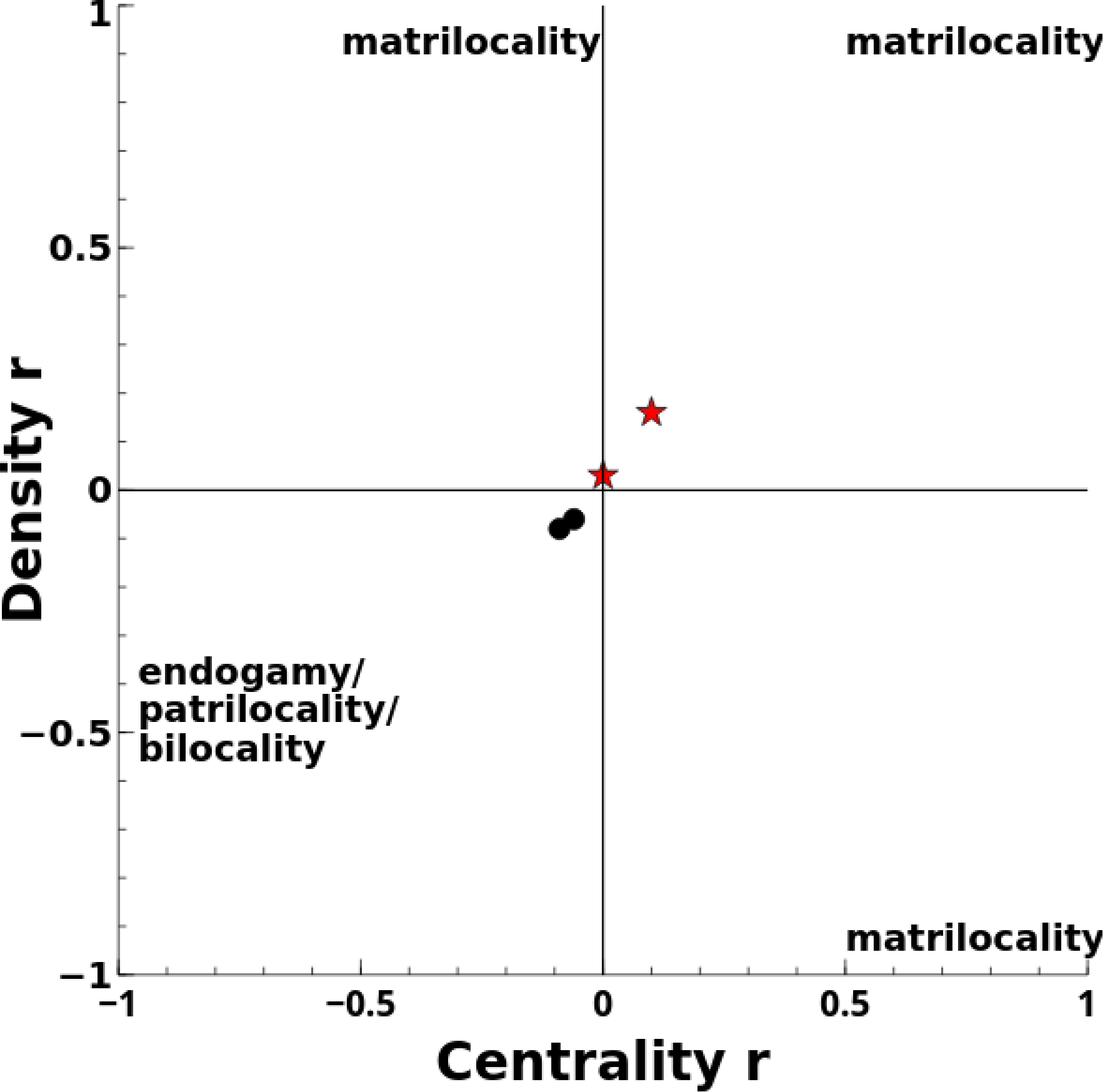
Scatterplot of centrality and density matrix correlations for haplogroups for Lokomotiv and Ust’Ida sites based on morphologic (black circles) and genetic (red stars) sex estimation methods

*Extinct hominins.* Lalueza-Fox and colleagues (111) studied 6 adult Neanderthal individuals, including 3 males and 3 females. The remains were discovered at El Sidron Cave in Asturias, Spain and dated to approximately 49,000 years before present. Lalueza-Fox and colleagues estimated male sex from a Y chromosome marker and female sex by the absence of this marker, combined with dental and morphologic criteria. They sequenced these individuals’ mtDNA at 11 phylogenetically significant nucleotide positions. Following Lalueza-Fox and colleagues, I treated the set of variants at these positions as a haplotype. The three males and one of the females had the same haplotype, but each of the other two females had a unique haplotype. By my calculation, the corresponding centrality hypothesis correlations for haplotype and haplotype similarity are -.41 (95% CI:-.92-.60) and -.24 (95% CI:-.87-.68), respectively. The density hypothesis correlations for haplotype and haplotype similarity are -1.00 (95% CI:-.98-.23) and -.87 (95% CI:-.99--.18), respectively. These results indicate a pattern consistent with endogamy, patrilocality, or bilocality.

Lalueza-Fox and colleagues (111) and Rosas and colleagues (112) noted that all of the individuals’ skeletal remains show evidence of human-induced modifications which they interpreted as cannibalism by fellow Neanderthals. In the other reports included in my review, authors did not describe evidence of cannibalism. Cannibalism in prehistoric humans and hominins may have sometimes been practiced on members of out-groups (113). Therefore, the El Sidron individuals may have hailed from groups that did not reside near the burial site, although their remains may be have been processed there by a group that was local to the area. This possibility renders any interpretation of post-marital residence patterns from the El Sidron remains problematic without isotope analyses and replication at other Neanderthal sites. Five years ago, Lalueza-Fox and colleagues (114) mentioned the prospect of strontium isotope analyses of the Neanderthal individuals. I have not found any report of such analyses in my search of the literature.

<u>Between site comparisons</u>. Table 8 shows sets of sites that were close to each other geographically and contemporaneous. The distances between sites listed in the table are geodesic distances (“as the crow flies”). Except for the Pyramid Lake-Stillwater Marsh pair of sites, all other sets of sites had partial or full water routes between them. Except for Aleutian set of sites, all other sets of sites had fairly direct, topographically benign overland routes. This table and Figure 11 also show the correlations that indicate the extent to which individuals at these sites have different mtDNA haplogroup distributions.

The summary estimate indicates a modest tendency toward female endogamy, very weakly in the direction of matrilocality (as opposed to endogamy overall), although these results are not statistically reliable. There is little genuine variation in the underlying estimates among the separate sets of sites (I^2^ =.19 for women and.00 for men). For each site, the correlation for women is more positive (greater distinctiveness between communities) than the correlation for men. There are data at the haplotype and haplotype similarity levels for the sites on Gotland, Sweden (71–75). For haplotypes, the inter-site correlations are.11 (95% CI, -.20-.41) for women and.40 (95% CI,.07-.74) for men, which together suggest endogamy. For haplotype similiarity, the inter-site correlations are.17 (95% CI, -.04-.37) and -.20 (95% CI, -.59-.19) for men, suggesting matrilocality. The haplogroup level correlations also suggest matrilocality (Table 8 and Figure 12).

In addition, Johnson (85) presented mtDNA results for individuals at two sites (Llano Seco [Table 4] and Wurlitzer [CA-BUT-294], 4430-1132 BP) approximately 32 kilometers apart, both in the northeastern Sacramento Valley near Chico, California, USA. No major topographical obstacles exist between them. Demographic data were available only for three adult individuals (2 male, 1 female) from the Wurlitzer site (CA-BUT-294), thus precluding a formal statistical analysis. The two Llano Seco men and the two Wurlitzer men had the same haplogroup (B). The Wurlitzer woman, however, had a different haplogroup (C) than each of the four Llano Seco women (three who had haplogroup D and one who had haplogroup B). These distributions are consistent with matrilocality.

**Table 8.**
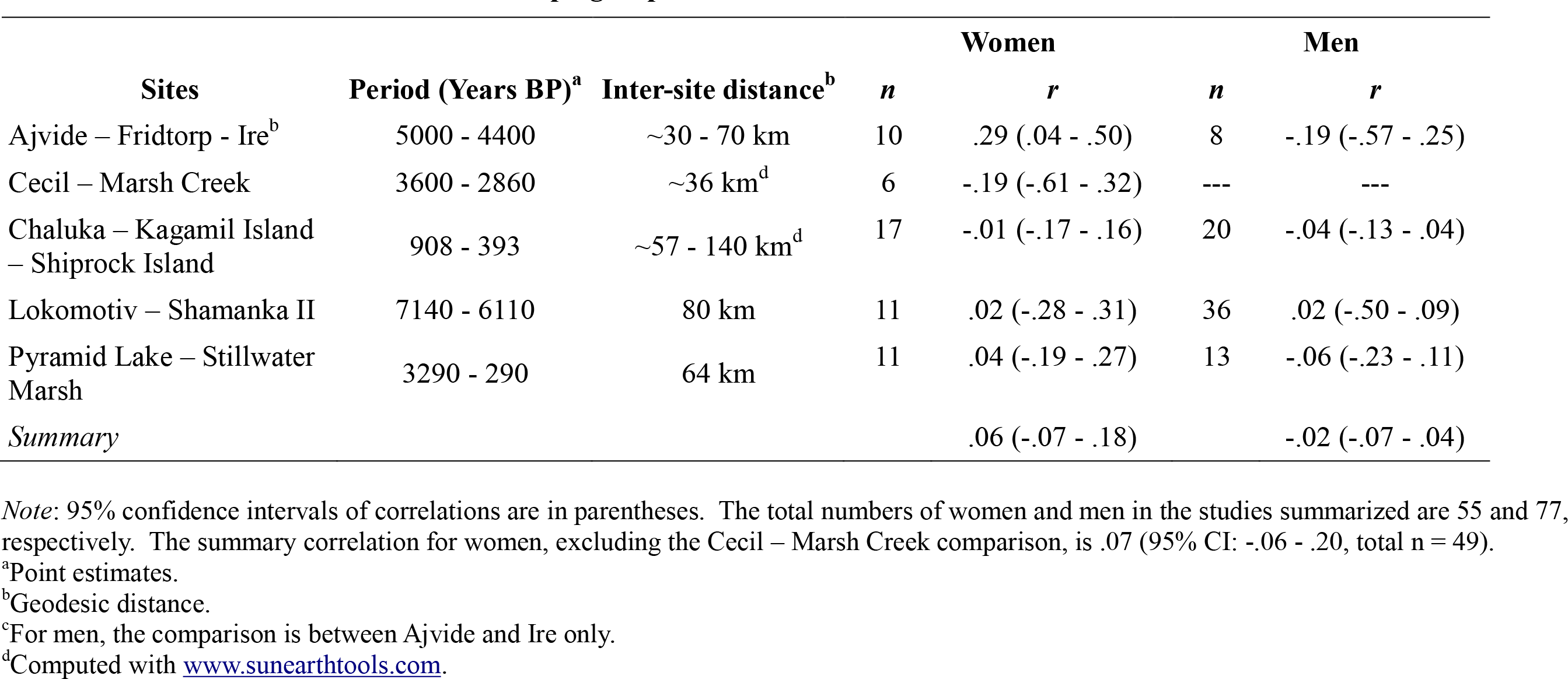
Inter-site matrix correlations for haplogroups

| Sites | Period (Years BP)<sup>a</sup> | Inter-site distance<sup>b</sup> | *n* | Women | *n* | Men |
| --- | --- | --- | --- | --- | --- | --- |
|  |  |  |  | *r* |  | *r* |
| Ajvide – Fridtorp - Ire<sup>b</sup> | 5000 - 4400 | ~30 - 70 km | 10 | .29 (.04 - .50) | 8 | -.19 (-.57 - .25) |
| Cecil – Marsh Creek | 3600 - 2860 | ~36 km<sup>d</sup> | 7 | -.24 (-.65 - .29) | --- | --- |
| Chaluka – Kagamil Island<br>– Shiprock Island | 908 - 393 | ~57 - 140 km<sup>d</sup> | 17 | -.01 (-.17 - .16) | 20 | -.04 (-.13 - .04) |
| Lokomotiv – Shamanka II | 7140 - 6110 | 80 km | 11 | .02 (-.28 - .31) | 36 | .02 (-.50 - .09) |
| Pyramid Lake – Stillwater<br>Marsh | 3290 - 290 | 64 km | 11 | .04 (-.19 - .27) | 13 | -.06 (-.23 - .11) |
| *Summary* |  |  |  | .06 (-.07 - .19) |  | -.02 (-.07 - .04) |

*Note:* 95% confidence intervals of correlations are in parentheses. The total numbers of women and men in the studies summarized are 56 and 77, respectively. The summary correlation for women, excluding the Cecil – Marsh Creek comparison, is .07 (95% CI: -.06 - .20, total *n* = 49).
<sup>a</sup>Point estimates.
<sup>b</sup>Geodesic distance.
<sup>c</sup>For men, the comparison is between Ajvide and Ire only.
<sup>d</sup>Computed with www.sunearthtools.com.

**Figure 12.**
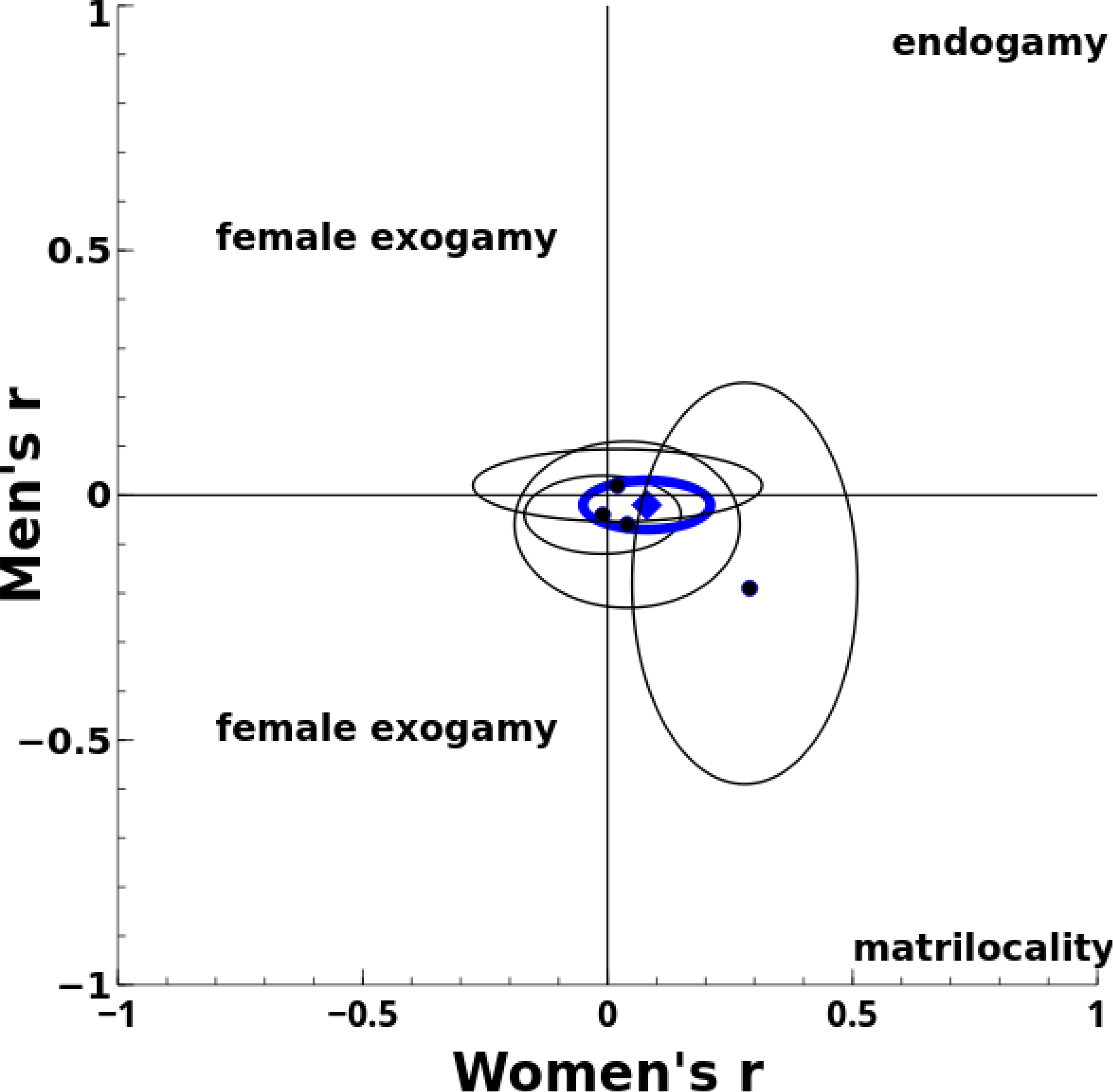
Scatterplot of women’s and men’s inter-site matrix correlations, including 95% confidence ellipses and summary estimate (blue, bold)

## Ethnographic Accounts of Burial Practices in Modern Hunter-Gatherers

### Methods

To evaluate my assumptions about what prehistoric hunter-gathers’ burial sites represent, I conducted a search of ethnographies of modern hunter-gatherers. I searched the eHRAF World Cultures database (http://ehrafworldcultures.yale.edu), created and curated by the Human Relations Area Files. The overall database includes 290 cultures and approximately 600,000 document pages. I used the “burial practices and funerals (764)” and “special burial practices and funerals (766)” subject terms for my search and inspected results for hunter-gatherer cultures, focusing especially on accounts of reported pre-contact behavior. If my assumptions about what prehistoric burial sites represent (i.e., buried individuals are representative of the burying community, a burial site corresponds to a single community over time, buried individuals lived as adults in the burying community, and outsiders killed in conflict are not in a community’s burial site) tend to hold for modern hunter-gatherers, then they might be reasonable assumptions for prehistoric hunter-gatherers in the absence of other information. While modern hunter-gatherers cannot necessarily serve as analogs of prehistoric hunter-gatherers, if my assumptions are inconsistent with modern hunter-gatherer behavior, it would not be sensible to extend these assumptions to prehistoric hunter-gatherers.

### Results

My search returned 3,330 paragraphs from 351 documents on 46 cultures. The relevant excerpts from these documents and my summaries of them are in supplementary files linked to this article (“eHRAFBurialPracticesExcerpts.pdf” and “Evaluation of eHRAF burial practice excerpts.pdf”).

Each of the ethnographic accounts suffers from major methodological problems. Statements about burial practices are not based on systematic data collection or analysis. Some statements are accompanied by one or a few anecdotal observations, but they often include no substantiating information. In many sources, it is not clear whether the ethnographer’s statements are based on his or her own experiences or those reported by other authors. Furthermore, ethnographers’ descriptions of aboriginal burial practices for a particular culture sometimes are inconsistent with each other. Consequently, I considered a practice to have been present if it is mentioned in most ethnographic accounts for a particular culture.

The ethnographic evidence is consistent with the assumption that individuals buried at group burial sites are generally representative of the burying community. The two main exceptions are the Alutiik (among whom high status persons, and sometimes their slaves, were buried at locations different from others) and Chinookans of the Lower Columbia River (among whom only slaves were buried in the ground). The Ainu represent a minor exception (persons who died on the water or were killed by bears were buried in different locations from other persons).

No ethnographic account contradicts the assumption that a group burial site corresponds to a single community over time. I found no reports of multiple communities sharing a burial site, or different communities through time using the same site.

The ethnographic evidence also shows that adults buried in cemeteries or sites of multiple burials were residents in the burying community. There are no reports of a community burying enemies killed during hostilities with the community’s own dead. Rather, the accounts indicated that the communities of the fallen attempted to recover all or part of the remains. There are also no reports of an immigrant individual being returned to his or her natal community for burial.

In 22 of the 45 cultures, burial practices would likely not result in archaeological detection of group burial sites after hundreds of years or more. In many of these cultures, corpses were buried in trees, on scaffolds, or on the surface at locations that were not spatially clustered with other similar burials. Scavenging animals probably would disperse human remains from such burial locations, and some ethnographers reported observations consistent with such dispersal. Burials or deposition of corpses in huts/lodges/tipis/igloos would also not produce archaeologically detectable group burial sites when the community abandoned camp after a death. Similarly, subsurface burials at unique locations and subsequent abandonment of camp would not result in group burial sites.

Cremation was practiced by four additional cultures (Klamath, Pomo, Yaghan, and Yokuts). Strontium isotope analysis of tooth enamel is usually impossible with cremated remains (115). Likewise, mtDNA usually cannot be accurately recovered from burned bone (116).

In four other cultures (Nuu-chah-nulth, Nuxalk, Quinalt, and Tlingit), burial practices might sometimes have produced archaeologically detectable group burial sites. In these cultures, tree, surface, and/or scaffold burials were typically done at spatially clustered group sites. Even though scavengers and surface geological processes might disperse remains at such locations, accumulations of remains over time or natural, sudden sedimentary covering of a location (e.g., from landslides, floods, or volcanic eruptions) could preserve the group nature of the burials. Also, these cultures sometimes used group cave burials, which likely produced archaeologically detectable group burial sites.

I found other relevant patterns in the ethnographic accounts. There were no reports of corpses being transported more than a few kilometers (usually they were moved very little, if at all). Most transports of more than a few hundred meters involved watercraft for much of the trip. There were some reports, however, of cremated or defleshed bones being transported significant distances, often involving conveyance by horse (a mode of transport not available to most prehistoric hunter-gatherers). In addition, there were no consistent indications that nomadic communities used sites of multiple burials or cemeteries.

## Discussion

### Summary of results

I summarized the evidence on post-marital residence patterns in prehistoric hunter-gathers from studies of strontium isotopes and mtDNA in human remains. The archaeologic sites included in my review represent every inhabited continent except Australia, and their dates span almost the last 10,000 years. As a whole, the studies I included have some features that increase confidence in the results. Many independent teams of authors, trained and based in several different countries across the world, produced the included reports. Authors and museums either published or made available the raw data underlying included reports. Moreover, publication bias was unlikely as very few authors even mentioned post-marital residence patterns in their reports.

In the strontium isotope studies, most adults of both sexes were local, and women were slightly more likely to be local than men. Within sites, women and men had similar mtDNA distributions. Women in neighboring, contemporaneous communities had somewhat distinct mtDNA distributions from each other, while the mtDNA distributions of men in the different communities were less distinguishable. Taken together, these results indicate the burial communities were mostly endogamous and that exogamous marriages strained toward matrilocality.

The very limited research on extinct hominins’ residence patterns seems to be consistent with these findings. In my re-evaluation of the evidence on extinct hominins, I found that the data actually pointed to endogamy in an isotope study of australopithecines, and endogamy, patrilocality, or bilocality (with no objective way to differentiate among these patterns) in an mtDNA study of cannibalized Neanderthal remains.

In a complementary systematic review, I found that there was only a moderate correspondence between morphologic and genetic estimates of sex in adult prehistoric hunter-gatherer individuals. When I corrected for this measurement error or relied on genetic estimates of sex only, the post-marital residence pattern results shifted in the direction of greater matrilocal tendencies.

As shown in my review of ethnographic accounts in the eHRAF World Cultures database, modern hunter-gatherers’ burial practices are consistent with my assumptions about what prehistoric hunter-gatherer burial sites represent. By these accounts, individuals buried at group burial sites were generally representative of the burying community, particular burial sites corresponded to single communities over time, and adults in group burial sites were residents of the burying community.

### Limitations

#### Overall

My literature searches were not focused specifically on strontium and mtDNA studies, and I may have missed some relevant reports because of this. While completing this article, I conducted some abbreviated searches with these more specific terms and found no additional pertinent reports.

Some of the hunter-gatherer societies included in this review may have been in contact, or even symbiotic relationships, with societies with other modes of subsistence (117,118). Such contacts could have altered hunter-gatherers’ post-marital residence patterns from their ancestral state.

Slavery, if present in a prehistoric hunter-gatherer society, could cloud interpretations of post-marital residence patterns inferred from strontium isotope and mtDNA studies. If individuals from one community captured individuals from another community and enslaved them and if deceased slaves were buried at the same sites as others in the capturing community, then genetic heterogeneity between communities would be reduced and immigrant isotope signatures and mtDNA would be introduced to the capturing community. The effects of slavery on mtDNA distributions would be multiplied if outsider women were enslaved and they reproduced with men from the capturing community. Slavery would dilute any observable tendencies toward endogamy and matrilocality (the latter in the case of imported female slaves).

To the extent that communities sometimes merged with each other, or incorporated immigrant residents apart from marriage, it would mask any tendency toward endogamy (in strontium isotope and mtDNA data) or matrilocality (in mtDNA data).

Most sites in my review span long periods, typically a few to many hundreds of years. Variation in post-marital residence patterns at a site over time would tend to obscure any prevailing pattern in a long period analyzed as a whole.

Prehistoric human remains do not indicate individuals’ marital statuses. Undoubtedly, I included some individuals who had never married, which would add noise to any signal of post-marital residence patterns. Sex differences in age at marriage (with men tending to be older) would introduce bias against detecting matrilocal tendencies if the community practiced any degree of male exogamy.

Authors typically did not describe in detail their procedures for estimating sex and age. The relatively low reliability I found in such classifications may be due not only to incomplete and damaged remains, but also observer error in morphologic assessments. For instance, when pairs of observers classified the sex of individuals from intact skeletal materials, they displayed only moderate agreement for crania (119,120) and somewhat higher agreement for pelvises (121).

The individuals included in my review are likely vastly outnumbered by the individuals on whom relevant data are missing. I excluded individuals for whom sex and age data were unavailable. Often, authors included only some individuals from a burial site in their studies, because of unexcavated remains, insufficient recovered remains, mtDNA too scarce to be amplified, lack of molar teeth, or other reasons. Burial sites also may represent only some of the locations a community used for burial. These missing data problems probably did not bias my results, but they decreased sample sizes. In addition, a few authors did not respond to or refused requests to share data and reports.

Although I summarized many studies, most had small samples, primarily due to the missing data problems. Consequently, the confidence intervals around my summary estimates are fairly wide and include tendencies toward opposite post-marital residence patterns (e.g., the confidence limits around strontium isotope estimates of weak matrilocal tendencies include a range from weak patrilocal tendencies to moderate matrilocality). As with nearly all systematic reviews, my results and conclusions are provisional and await refinement by the addition of new evidence. Nonethless, based on the available data, moderate to strong exogamy and moderate to strong patrilocality can be rejected as post-marital residence patterns in prehistoric hunter-gatherers.

#### Strontium isotope studies

The geographic scope of a particular local isotope ratio range has a significant influence on the classification of individuals as locals or immigrants. If the isotopic signature varies little over a large geographic area and multiple concurrent communities were present in that area, individuals raised in different communities could have similar isotope ratios. This means that some individuals classified as locals could actually have been immigrants to the communities in which they were buried.

Conversely, members of a community may have ranged more widely than the geographic scope of the measured local isotope ratios. That is, the sampled locations for modern flora, sediments, or water, or the home ranges of archaeologic fauna may not have been as extensive as the territory a given set of prehistoric hunter-gatherers traversed. The measured local isotope ratio range could, as a result, be biased from this mismatch. This is particularly so when the sampling did not extend beyond the site itself. Even sedentary hunter-gatherers in rich environments likely covered relatively large areas for subsistence, probably varying by season or multiple year periods. If their territories were larger than those for the sampled isotope ratios, some locals may have been misclassified as immigrants.

It is not possible to assess mobility in childhood for strontium isotope studies in which only one tooth per individual was available or tested. Such mobility, if present, would prevent a clear interpretation of post-marital residence patterns.

#### mtDNA studies

Contamination is a constant and universal threat in ancient DNA studies (122). Contamination from one sample to another (e.g., through aerosols created during routine lab work and the resulting tainted equipment) is a particular concern for my review, as the genetic relatedness between individuals (samples) was the dependent variable. Such processes could even create, artifactually, the genetic differences between contemporaneous neighboring communities, as the individuals from the different sites were or may have been analyzed in different labs or in the same lab at different times.

Some natural factors could make post-marital residence patterns difficult to discern from mtDNA distributions. If two communities exchange female mates (female exogamy) without respect to haplotype, even at low levels that could even be much lower than for the exchange of male mates (male exogamy), over time they will tend to have similar mtDNA distributions in both sexes due to simple probabilistic gene flow. That is, unless matrilocality or endogamy is complete, with no women moving after marriage, repeated instances of female exogamy eventually lead to similar mtDNA distributions. Likewise, neighboring prehistoric hunter-gatherer communities often may have been genetically related to each other, such as when they both descended from the same ancestral group. This could produce a baseline of genetic similarity between the communities, reducing the chances of detecting distinctive genetic signatures of different, nearby communities, especially in small samples.

The between-site mtDNA comparisons are based on the assumption that the different sites represent different communities. If, however, a pair of sites belonged to the same community (e.g., at different locations within their range), then the comparisons would not be meaningful.

Furthermore, prehistoric hunter-gatherer populations tended to have no to very small population growth (123,124). In such circumstances, most women would, on average, propagate their mtDNA only a few generations forward. These mitochrondial DNA matrilines typically would be slender and include a handful or fewer adult individuals. The high haplotypic diversity in the studies I reviewed is consistent with narrow and short mtDNA matrilines. The sparsity of the haplotype data might have attenuated any patterns that might have present in these communities.

#### eHRAF World Cultures

This database is not a complete inventory of ethnographies of modern hunter-gatherers, and therefore it is possible that there are some cultures with burial practices that violate the assumptions on which my review is based. Also, the many significant methodological shortcomings of the non-systematic ethnographies included in the database make it an imperfect resource. However, despite the lack of methodological rigor, it seems very unlikely ethnographers were biased in a way that aligned so neatly with my assumptions.

### Generalizability

The results from the strontium isotope and mtDNA studies were consistent with each other. However, there were no sites from Australia, South Asia, or Southeastern Asia included in my review. Moreover, my summaries, at best, can only be generalized to prehistoric hunter-gatherers who bury their dead in common locations. My review of the eHRAF World Cultures ethnographic data indicated that hunter-gatherers living in arid, tropical, or cold environments have burial practices that are unlikely to result in archaeological burial sites. Authors of the included reports and other researchers interpreted the archaeological evidence from their sites as indicating the communities were sedentary or semi-sedentary (when they offered interpretations of mobility patterns). The connection between sedentism and common burial sites was also apparent in the eHRAF World Cultures ethnographic data on hunter-gatherers. Taken together, these observations mean that my summaries might not generalize to nomadic hunter-gatherers.

Although many anthropologists assume that hunter-gatherers were exclusively nomadic until the Holocene, the evidence indicates and implies that hunter-gatherer sedentism, especially that associated with marine, estuarine, or lacustrine environments (as in most of the sites included in my review), may be represented throughout human evolutionary history (125). Thus, both nomadic and sedentary subsistence patterns may be part of the ancient human repertoire. Indeed, sites of many modern human burials occurring over short periods of time date to at least 90,000 years ago (126). Sites of funerary caching of many (>12) corpses, such as surface abandonment at a particular location, extend back to the Pliocene (126,127). Rowley-Conwy (125) hypothesized that the lack of excavated sites showing clear evidence of sedentary human communities in the Pleistocene is due to sea level changes. That is,Pleistocene hunter-gatherers primarily exploited marine and estuarine habitats and most such areas in the Pleistocene are now underwater and undetected.

### Evolution of post-marital residence patterns

Even though my summaries of the evidence are far from definitive, I nonetheless speculate on proximate and ultimate causes of the post-marital residence patterns I found, following Twain’s apt description of scientific practice. With respect to proximate causes, if prehistoric hunter-gatherers maintained low population densities with widely separated and non-overlapping ranges, endogamy might the natural result, as Froment (128) postulated for nomadic hunter-gatherers. Contact between communities may have been too infrequent and difficult to sustain much exogamy. While such extreme circumstances may have held at many points and places in human evolution, many of the sites included in my review seem not to have been so severely isolated.

The sexual division of labor might also have been a proximate driver toward matrilocal tendencies. As the primary hunters and fishers, even in sedentary communities, men likely had long and regular absences from home base, as they sought game, acquired other resources from distant locations, and engaged in trade. Men probably pushed geographic frontiers and encountered other communities more often than women did. With all other factors equal, it might be more likely that a community would accept a strange or relatively unfamiliar individual-a man-as a new resident than it would be for a community to let one of its members-a woman-leave with a strange or relatively unfamiliar individual.

Ultimate causes of post-marital residence patterns can be identified by viewing such patterns as the outcomes of long-term mating preferences. Regarding sex-biased dispersal patterns in mammals generally, Handley and Perrin (23) emphasized that “mating systems are thus a priori expected to affect dispersal patterns” (p. 1560). A hallmark of long-term mating in sexual species in which fathers invest in their offspring, such as humans, is that females choose males (129). Women, as with many female mammals, by their biological nature invest more in offspring than men. Hence, in the absence of other factors, women hold more leverage than men in setting many of the conditions for marriage. Men compete for women, who are the scarce reproductive resource. One of the key criteria women use in choosing a long-term mate is his ability to invest in children, which is typically expressed as wealth or wealth potential (130).

In hunter-gatherers, a man’s wealth is primarily stored in his person-skills, knowledge, personality traits, relationships, and physical strength, speed, agility, and endurance. Indeed, ethnographic data from a broad sample of modern hunter-gatherer societies showed that hunter-gatherers’ most preferred traits in a son-in-law were that he was a good hunter, hard worker, good provider, and from a good family (131). In this sample, parents and other kin were the primary decision makers for marriages (131).

It may be advantageous for a woman’s kin to keep her close after she marries, so they can invest more intensively in her offspring than if she lived in another community. Biological kin tend to invest more in a woman’s children than a man’s children (132,133), as the woman’s children are definitely her own, while paternity typically is somewhat uncertain. Just two years after Darwin published *On the Origin of Species,* Bachofen (134) gave similar reasons for proposing that matriliny was the ancestral human system of descent.

Of all post-marital residence patterns, endogamy might tend to be the most advantageous to children, parents, and their kin by increasing the inclusive fitness of all. Endogamy allows the best opportunity for the woman and her kin to evaluate potential mates over a long period of time in the same community. This is important, because the traits most relevant to a potential husband’s fitness (skills, knowledge, personality, etc.) can only be assessed reliably over time. Similarly, endogamy enables a man and his kin to judge more effectively some of those traits that men prefer in a long-term mate, such as chastity (130).

Furthermore, under endogamy, both wife’s and husband’s kin can invest intensively in the couple’s offspring. Indeed, endogamy and the unique human brand of cooperative breeding (11) may have coevolved. In endogamous communities, even if a wife cuckolds her husband, he still tends to have some genetic relationship to her offspring. Also, to the extent that prehistoric hunter-gatherers suffered mortality before middle and old age (11), endogamy would maximize the number of kin immediately available to help with child rearing and support. Still, endogamy and/or matrilocality almost certainly were not complete, because the moderate to high mitochondrial diversity at nearly every site in my review was probably due, in large part, to some degree of past female exogamy, possibly at very low levels for many millennia.

There is some evidence that endogamy in small communities enhances reproductive fitness. In scores of modern hunter-gatherer, horticultural, agricultural, pastoral, and industrial societies across all major cultural regions of the world, a married couple’s fertility declined with decreasing genetic relatedness (135–142). A couple’s number of surviving children typically peaked on average for levels of relatedness equal to second, third, or fourth cousins, and then was markedly less at lower levels of relatedness. Helgason and colleagues (137) further found in their analyses of all marriages in Iceland from 1800 to 1965 that the number of children who reproduced and the number of grandchildren also peaked at levels of genetic relatedness between spouses equal to third and fourth cousins. In light of this evidence, Fox (143) reformulated Tylor’s “marry out or die out!” commandment to “marry IN or die out!” Fox also linked this dynamic to the fissioning of communities in small-scale societies once the average genetic relatedness falls below a threshold of about fourth cousins. In prehistory, a moderately to strongly exogamous marriage system that paired unrelated persons from separate communities probably was not an evolutionary stable state.

After hunter-gathers have substantial contact with or transition into horticultural, pastoral, or agricultural societies, men can usually store wealth outside of their persons in land, livestock, large physical objects, and/or geographically fixed resources that are not practically portable or not portable at all. To the extent that wealth is a critical factor in selecting a husband, these methods of storing wealth may shift the leverage in marriage to men and lead to the dominance of patrilocality (144–146). In fact, in 1881, Morgan (147) noted that in human evolution, once people stored wealth in property, descent systems became patrilineal.

### Implications

As I noted in the introduction, for more than 150 years anthropologists have generally presumed that prehistoric hunter-gatherers were exogamous, debating only the type of exogamy they practiced. I was heavily influenced by this bias in the literature and initially geared my review accordingly. Yet endogamy could have many biological and social implications, and might even be necessary to account for key aspects of human health and behavior. I focus my conjecture on just two:sexually transmitted diseases and altruism.

#### Sexually transmitted diseases

Endogamy in small populations of prehistoric hunter-gatherers likely inhibited the evolution of bacterial sexually transmitted diseases (STD) by preventing sexual connections between communities and thus producing highly fragmented sexual networks. Froment (128) speculated similarly with respect to nomadic prehistoric hunter-gatherers, although I posit that these circumstances held for sedentary hunter-gatherers as well. Indeed, both syphilis and gonorrhea seem to be relatively recently evolved human pathogens. Osteologic diagnosis of syphilis can only be made with confidence for congenital syphilis in children, as the signs of syphilis and two other treponemal (but non-venereal) infections, yaws and bejel, overlap too much in adult remains to differentiate these diagnoses (148). Nearly all of the earliest cases of possible and probable congenital syphilis, based on skeletal and dental evidence, identified to date were in agricultural societies in the Americas and Europe and dated within the last 2500 years (149–152). Pruemers and colleagues (153) detected DNA of the bacterium causing venereal syphilis in individuals found at a pre-Columbian (500-1400 AD) agricultural site in Bolivia and whose skeletal remains showed signs pathognomic for treponemal infection. If syphilis has mutation rates similar to other bacteria, then the phylogenetic relationships among geographically disparate modern syphilis strains imply that syphilis diverged from other treponemal pathogens more than 500 years ago and perhaps as long as several thousand years ago (154). Based on the historical evidence, gonorrhea seems to have first evolved into a sexually transmitted infection in medieval Europe (155).

Over the many hundred thousands of years of human evolution, it appears that bacterial STD were only able to gain a foothold in humans within the last 10,000 years, once some people adopted an agricultural mode of subsistence, urban settlement, and exogamy. These circumstances enabled large connected populations of hosts in which a bacterial pathogen could circulate and allowed subpopulations of individuals with dense sexual networks to form who serve as a reservoir for infection (156,157). However, viral STD, which tend to have much longer periods of communicability, would not necessarily have been inhibited by endogamy to the same degree as bacterial STD.

#### Altruism

Kin selection (favoring the reproductive success of one’s biological kin, even at cost to self) and reciprocal altruism (behavior that increases another’s fitness at temporary cost to self and is likely to be reciprocated in the future) underlie the evolution of altruism (158–160). Endogamy produces the circumstances by which kin selection and reciprocal altruism can operate on a regular basis. On one hand, individuals in small endogamous communities tend to have high average genetic relatedness to each other. My summaries suggest that prehistoric hunter-gatherers’ kin and fellow community members were almost coterminous. On the other hand, endogamous communities satisfy two preconditions for the evolution of reciprocal altruism:low dispersal rates and small, mutually dependent, stable social groups (160). Exogamy would prevent both kin selection and reciprocal altruism from operating pervasively. The combination of endogamy and cooperative breeding (11) could result in the evolution of generalized human dispositions toward sharing and generosity. Moreover, endogamy, especially for men, might also have been necessary for the evolution of psychological adaptations that leave people susceptible to such group-level phenomena as warfare, ethnocentrism, and xenophobia.

### Future research

I suspect there are many collections of prehistoric hunter-gatherer remains that would be suitable for strontium isotope and/or mtDNA analyses. Such work would help to narrow confidence intervals of summary estimates of post-marital residence patterns, and possibly fill geographic gaps and extend the time range of sites studied.

Ideal study designs would involve both strontium isotope and mtDNA methods applied to the same samples of individuals drawn from adjacent, contemporaneous communities. Researchers studying sites in the Lake Baikal and California Delta regions have done this (35,41,44–47,62,78,91–93,99), although the strontium isotope results have not always yielded clear interpretations, likely due to the mobility of some communities. Comparisons of men’s Y-chromosome haplogroup/haplotype distributions in such concurrent, neighboring communities would be valuable. With the same hypothesis structure I used for between-site comparisons, a positive matrix correlation for Y-chromosome data would indicate endogamy or patrilocality, and a negative correlation would indicate male exogamy.

It is important to measure strontium isotope ratios of first, second, and third molars in the same individuals, if possible, for assessing the mobility of the community as a whole. If a meaningful proportion (say > 15%) of adults have both local and non-local signatures in their molar enamel, it suggests that the burial community’s range is larger than that reflected in the isotope measurements of the “local” area and that the community may have had other burial sites. Such mobility in childhood could also reflect fusion of different communities. In either case, it may be difficult to interpret post-marital residence patterns objectively. Similarly, it is also critical to define local isotopic ranges based on sampling procedures and materials that correspond geographically to likely hunter-gatherer ranges.

Genetic measurement of sex would also be desirable in future research. When genetic measurement is not feasible, reliability of osteological/dental measurements could be improved with multiple independent observers. It would be interesting to assess whether combining observers’ judgments (161) improves accuracy as determined by a genetic measure.

My project was feasible because most authors reported the critical data in their publications. I made significant efforts to obtain additional information from authors who did not publish all necessary information. This experience underlines the importance of depositing data in public archives, especially in cases where the full data are not published in research reports. Without such practices, researchers effectively reduce the scientific value of their work.

## Acknowledgments

Many authors and other professionals kindly provided unpublished data, including the following:Noburu Adachi; Murilo Bastos; Clio Der Sarkissian; Glen Doran; Chris Dudar of the Repatriation Office of the National Museum of Natural History; Elizabeth Guerra of the University of California, Davis, Department of Anthropology Museum; Fredrik Hallgren; Johannes Krause; John Krigbaum; Rachel Malloy of the Nevada State Museum; Helena Malmstrom; Naomi Martisius; Vyacheslav Moiseyev; Amiee Potter; David Reich; Ken-ichi Shinoda; Beth Shook; Silvia Smith; Jan Stora; and Andrzej Weber. I am especially grateful to Keith Johnson for locating, recording, and sending unpublished data in difficult circumstances. Eric Bartelink, Alex Bentley, and Helena Malmstrom gave helpful references, and Sachiko Kato generously facilitated contact with some authors. John M. Roberts, Jr. located and retrieved a source. I also had helpful discussions with him about statistical issues. Linda Dick-Bissonnette, Carol Ember, Barbara Leigh, John Potterat, Dwight Read, Kim Romney, John Roberts, Jr., and Matt Sponheimer gave helpful comments on an earlier version of this article. I am responsible for all remaining errors and omissions. I received no funding for this research. All supplementary files for this article are also available at https://osf.io/ea3nm.

